# Folic acid polyglutamylation triggers MAP2K5 signaling to induce adipocyte lipid catabolism

**DOI:** 10.64898/2026.09.15.751849

**Authors:** Woo Sung Kim, Rebecca Danner, Juyeong Cho, Jonathan Williams, Luiz Lopez, Yutong Bao, Julia K. Hansen, Elizabeth M. Poad, Haania Khan, Daehan Kim, Brian W. Parks, Jonathan G. Van Vranken, David A. Harris, Andrea Galmozzi, Mary Elizabeth Patti, Snehal N. Chaudhari

## Abstract

Adipocyte signaling adaptively responds to macronutrients, but how essential micronutrients impact lipid homeostasis remains poorly understood. Here, we demonstrate that folate polyglutamylation, the sequential conjugation of glutamate residues to folate, serves as a dynamic biochemical process regulating metabolic signaling. Using our folate metabolomics platform, we show that polyglutamylated folic acid accumulates in healthy adipose tissue, but is depleted in obesity across mice and humans, independent of circulating folate levels. Genetic ablation of the polyglutamylation enzyme folylpolyglutamate synthase (Fpgs) in adipocytes suppresses lipid catabolism to induce cell-autonomous lipid accumulation, operating independently of canonical adipogenesis or downstream one-carbon flux. Target-engagement proteomics identifies monoglutamylated folic acid as a Map2k5-interacting molecule that inhibits Map2k5 activity, whereas Fpgs-mediated folic acid polyglutamylation acts as a chemical switch that disrupts this interaction to promote lipolysis. *In vivo*, whole-body inhibition of Map2k5 or adipose-targeted genetic depletion of Fpgs increases body fat mass in the absence of dietary obesogenic triggers. Furthermore, single-cell and single-nuclear transcriptomic analyses establish the Fpgs-Map2k5 pathway as a core transcriptional signature of mouse and human adipose tissues. These findings uncover a non-canonical signaling role for folate that regulates adipocyte lipid homeostasis.

## INTRODUCTION

Our understanding of obesity has evolved from a disorder of excessive energy storage to a progressive failure of adipose tissue homeostasis^1^. At the core of this dysfunction is the expansion of adipocytes, characterized by an increase in adipocyte size due to excessive lipid accumulation^2^. Pathological adipocyte hypertrophy is a hallmark of obesity and serves as a primary driver of local insulin resistance and inflammation, eventually leading to deterioration of systemic metabolic health^3^. Obesity is thus a leading risk factor for multifactorial metabolic diseases such as type 2 diabetes, cardiovascular diseases, and cancers, affecting over a billion people worldwide^4^.

Dietary nutrient availability is a major determinant of adipocyte expansion and hypertrophic remodeling^5^. Historically, nutritional research in obesity has focused largely on the manipulation of macronutrient ratios^6-8^. While the metabolic consequences of varying carbohydrate, lipid, and protein intake on adipocyte hypertrophy have been extensively studied, micronutrients also participate in metabolic processes that could influence adipocyte function. Folate (vitamin B9) is an essential cofactor in one-carbon metabolism (OCM), supporting nucleotide biosynthesis, amino acid metabolism, and S-adenosylmethionine production for methylation reactions^9^. Beyond this canonical role in cellular growth and proliferation, folate has been linked to obesity for many decades. Studies examining the relationship between dietary folate intake and body weight have, however, yielded inconsistent results. While some rodent studies associate folate deficiencies with weight gain^10-12^, others associate excess folate intake with increased body weight^13^. Studies in humans similarly report conflicting associations between folate intake or circulating folate levels and body mass index (BMI)^14-16^. These observations suggest a disconnect between folate intake and obesity, and that systemic folate status is a poor proxy for its metabolic activity within adipose tissues. Such discrepancies underscore the necessity of shifting focus from dietary or circulating levels of folate to the intracellular mechanisms through which folate directly regulates adipocyte function and hypertrophic growth. Previous studies show that depletion of folate in growth media increases intracellular lipid levels and promotes hypertrophy *in vitro* in adipocytes derived from murine and avian models^10,17^. However, the molecular mechanisms underlying the role of folate in directly modulating adipocyte lipid storage are unknown.

To profile tissue resident folates, we previously charted a comprehensive atlas of folate metabolites across 37 mouse tissues, which revealed organ-specific variations in folate vitamer distribution (Figure 1A and B)^18^. We observed a striking heterogeneity in folate polyglutamylation (Figure 1C), a unique intracellular reaction wherein multiple glutamate residues are sequentially conjugated to folate vitamers. Contrary to its traditional view as a folate-retention mechanism regulated primarily by folate availability, our work revealed that folate polyglutamylation is governed by tissue-specific transcriptional programs controlling the opposing activities of folylpolyglutamate synthase (*Fpgs*) and γ-glutamyl hydrolase (*Ggh*) enzymes, which add and remove glutamate residues respectively. Adipose tissues, including brown adipose tissue (BAT) as well as inguinal, perigonadal, and mesenteric white adipose depots (iWAT, gWAT, and mWAT) are enriched in two major folate species: folic acid (FA) and 5-methyltetrahydrofolate (5-m-THF). Notably, we found that adipose tissue FA is exclusively polyglutamylated while 5-m-THF is predominantly monoglutamylated, revealing distinct polyglutamylation patterns among folate vitamers in adipose tissue^18^.

**Figure 1.**
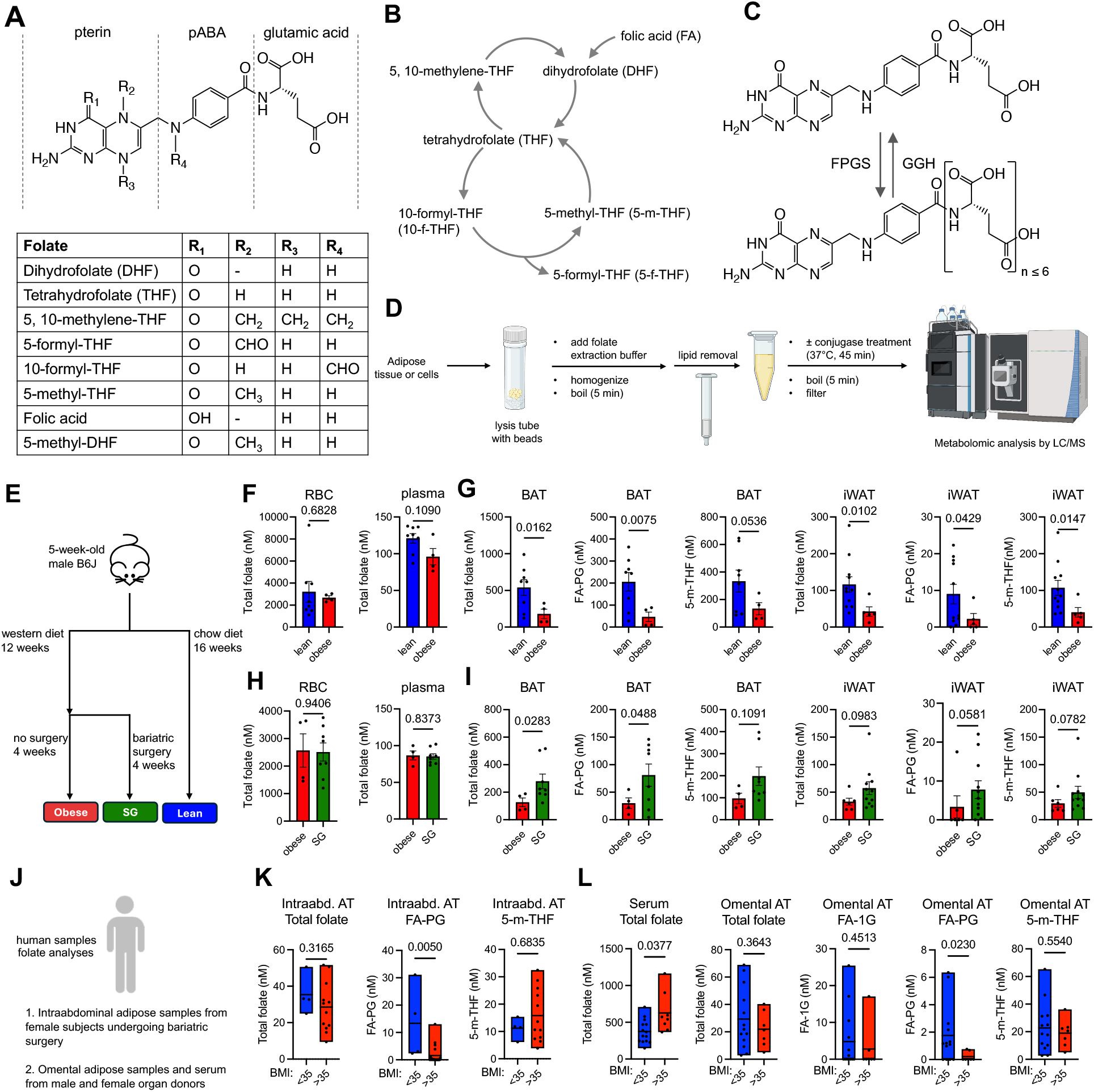
Adipose resident folates are depleted in obesity. (A) Structure of folate vitamers. (B), The folate one-carbon metabolism cycle. (C) Polyglutamylation of folic acid mediated by folylpolyglutamate synthase (FPGS) enzyme, and deglutamylation mediated by the γ-glutamyl hydrolase (GGH) enzyme. (D) Adipose folate metabolomics workflow showing extraction, solid phase extraction (SPE)- mediated lipid removal, and LC-MS analyses. (E) Schematic of mouse experiment and timelines. (F) Folate levels in red blood cells (RBC) and plasma from lean and obese mice (lean *n* = 8, obese *n* = 4, Mann-Whitney test for RBC, Welch’s t test for plasma). (G) Folate levels in brown adipose tissue (BAT) and inguinal white AT (iWAT) from lean and obese mice (BAT: lean *n* = 8, obese *n* = 4; iWAT: lean *n* = 11, obese *n* = 5, Welch’s t test). (H) Folate levels in RBC and plasma from obese and sleeve gastrectomy (SG) mice (obese *n* = 4, SG *n* = 8, Welch’s t test). (I) Folate levels in BAT and iWAT from obese and SG mice (BAT: obese *n* = 4, SG *n* = 8; iWAT: obese *n* = 6, SG *n* = 11, Welch’s t test for BAT FA-PG, Mann-Whitney test for all other comparisons). (J) Human cohorts analyzed in this study. (K) Folate levels in intraabdominal adipose tissue samples from female subjects undergoing bariatric surgery (BMI <35 *n* = 4, BMI >35 *n* = 12). l, Folate levels in omental adipose samples from the University of Wisconsin Health Human Organ Donor Program (BMI <35 *n* = 13, BMI >35 *n* = 7). Bar graphs are represented as mean ± SEM. Floating bars represent all data points with a line at mean. All data points represent biological replicates. Exact *P* values are shown in the graphs.

In this study, we investigated how adipose tissue-resident folates influence adipocyte lipid metabolism and hypertrophic growth. We uncover a previously unrecognized role for folate polyglutamylation in adipocyte metabolism and identify a mechanism through which the dietary form of folate regulates lipid homeostasis. These findings reveal an unexpected, non-canonical function for folate metabolism in adipose tissue and establish a direct link between folate processing and adipocyte metabolic function.

## RESULTS

### Adipose resident folates are depleted in obesity

While previous studies have quantified systemic folate levels in obesity across animal models and human cohorts, the specific folate profiles of adipose tissues have remained uncharacterized. This gap is primarily due to the severe analytical challenges in achieving robust recovery of highly polar folate metabolites from hydrophobic, lipid-rich adipose depots. Further, folates represent a chemically diverse class of metabolites with high structural similarities and a propensity to interconvert during sample processing (Figure 1A-C). This inherent instability, combined with the existence of multiple cellular redox states makes the precise profiling of individual folate species within tissues exceptionally challenging. To overcome these limitations, we developed an advanced liquid chromatography-mass spectrometry (LC-MS) folate metabolomics platform that enables the isolation and comprehensive profiling of chemically diverse folate molecules from a variety of complex biological matrices^18^. For adipose tissues specifically, our method utilizes an antioxidant-rich extraction buffer coupled with solid phase extraction (SPE)-mediated lipid removal step to achieve reproducible folate recovery (Figure 1D and Table S1).

To investigate whether adipose tissue folate abundance is altered in obesity, we quantified folate species in mice maintained on either a standard chow (lean) or western (obese) diet (Figure 1E). Folates in four distinct adipose tissues (brown, inguinal white, mesenteric white, and gonadal white adipose tissue; BAT, iWAT, mWAT, gWAT) and blood (both red blood cells and plasma) were quantified. Red blood cell (RBC) folate did not differ between lean and obese mice, and plasma folate showed only a modest, non-significant decrease in obesity^14^ (Figure 1F). In contrast, total folate abundance was markedly reduced in BAT, iWAT, gWAT and mWAT of obese mice compared to lean controls (Figure 1G and Figure S1A). Folic acid polyglutamate (FA-PG) and 5-methyl-tetrahydrofolate (5-m-THF), two major folate vitamers found in adipose tissues^18^, were both decreased across adipose depots in obesity (Figure 1G and Figure S1A).

To determine whether these folate metabolite shifts were reversible and independent of diet composition, we performed sleeve gastrectomy (SG) on western diet-fed obese mice to induce weight loss, while maintaining the western diet post-surgery (Figure 1E). Circulating folate concentrations remained unchanged following SG (Figure 1H). However, SG-mediated weight loss induced a 2-fold increase in total folate levels in BAT, iWAT, and mWAT, accompanied by a minor increase in gWAT (Figure 1I and Figure S1B). Consistently, folate levels in BAT, iWAT, and mWAT exhibited significant negative correlation with body weight, whereas plasma and gWAT displayed a non-significant, yet trending negative correlation (Figure S1C). In contrast, RBC folates were not correlated with body weight (Figure S1C). We next examined whether dynamic changes in body weight impact metabolic pathways downstream of folate by quantifying one-carbon metabolism (OCM) and energy metabolic intermediates in adipose depots. Targeted metabolite profiling revealed body weight-associated negative correlations that were primarily localized to BAT, with sparse correlations in WAT depots (Figure S1D). These findings together indicate that adipose folate levels are more sensitive to changes in body weight than systemic folate, without evidence of widespread metabolic alterations specifically in white adipose depots.

To test the clinical translation of our findings, we next evaluated whether the adipose-specific folate depletion observed in mice extends to human obesity, where adipose tissue folate profiles remain uncharacterized. We analyzed adipose tissue from two distinct human cohorts: (1). intraabdominal adipose samples from female subjects undergoing bariatric surgery, and (2) omental adipose samples from the University of Wisconsin Health Organ and Tissue Donation (UW-OTD) program (Figure 1J). We found that female subjects undergoing bariatric surgery with class 2 and 3 obesity (BMI > 35) exhibited a severe depletion of FA-PG in intra-abdominal adipose tissue compared to the leaner cohort, while 5-m-THF levels remain relatively unchanged (Figure 1K). A similar depletion of FA-PG in class 2 and 3 obesity was observed in omental adipose samples from human UW-OTD donors, even though serum folate levels were increased in these individuals (Figure 1L). Collectively, these data demonstrate an adipose tissue-specific regulation of folate metabolism in response to obesity in mice and humans independent of systemic folate availability.

### Folic acid polyglutamylation drives lipid catabolism to prevent adipocyte lipid accumulation

Given that both FA-PG and 5-m-THF levels were depleted in adipose tissues during obesity (Figure 1), we next investigated whether depletion of these folate vitamers influences lipid metabolism in adipocytes. We utilized the murine 3T3-L1 fibroblast cell line, a well-established *in vitro* model that recapitulates mature adipocyte morphology and metabolic pathways upon differentiation^19,20^. We confirmed that the baseline folate composition in 3T3-L1 cells mirrors *in vivo* adipose tissue, also exhibiting FA and 5-m-THF (Figure S2A and S2B). To impair intracellular folate polyglutamylation, we performed siRNA-mediated knockdown (kd) of *Fpgs* in 3T3-L1 cells, which reduced transcript levels by 50% (Figure S2A). *Fpgs* knockdown significantly suppressed folate polyglutamylation, yielding a significant 90% decrease in FA-PG and a 50% decrease in polyglutamylated 5-m-THF (Figure S2B). Strikingly, *Fpgs* deficiency triggered precocious lipid accumulation in preadipocytes, prior to the initiation of the standard differentiation program (Figure 2A). This elevation in intracellular lipid levels was maintained throughout differentiation and persisted in mature adipocytes (Figure 2B). Ultrastructural analysis via transmission electron microscopy confirmed an increased abundance of lipid droplets in *Fpgs* kd preadipocytes relative to controls (Figure 2C), supporting a role for folate polyglutamylation in limiting intracellular lipid storage. These results demonstrate that folate polyglutamylation influences basal lipid homeostasis across the adipocyte lineage that manifests in early progenitors and persists through maturity.

**Figure 2.**
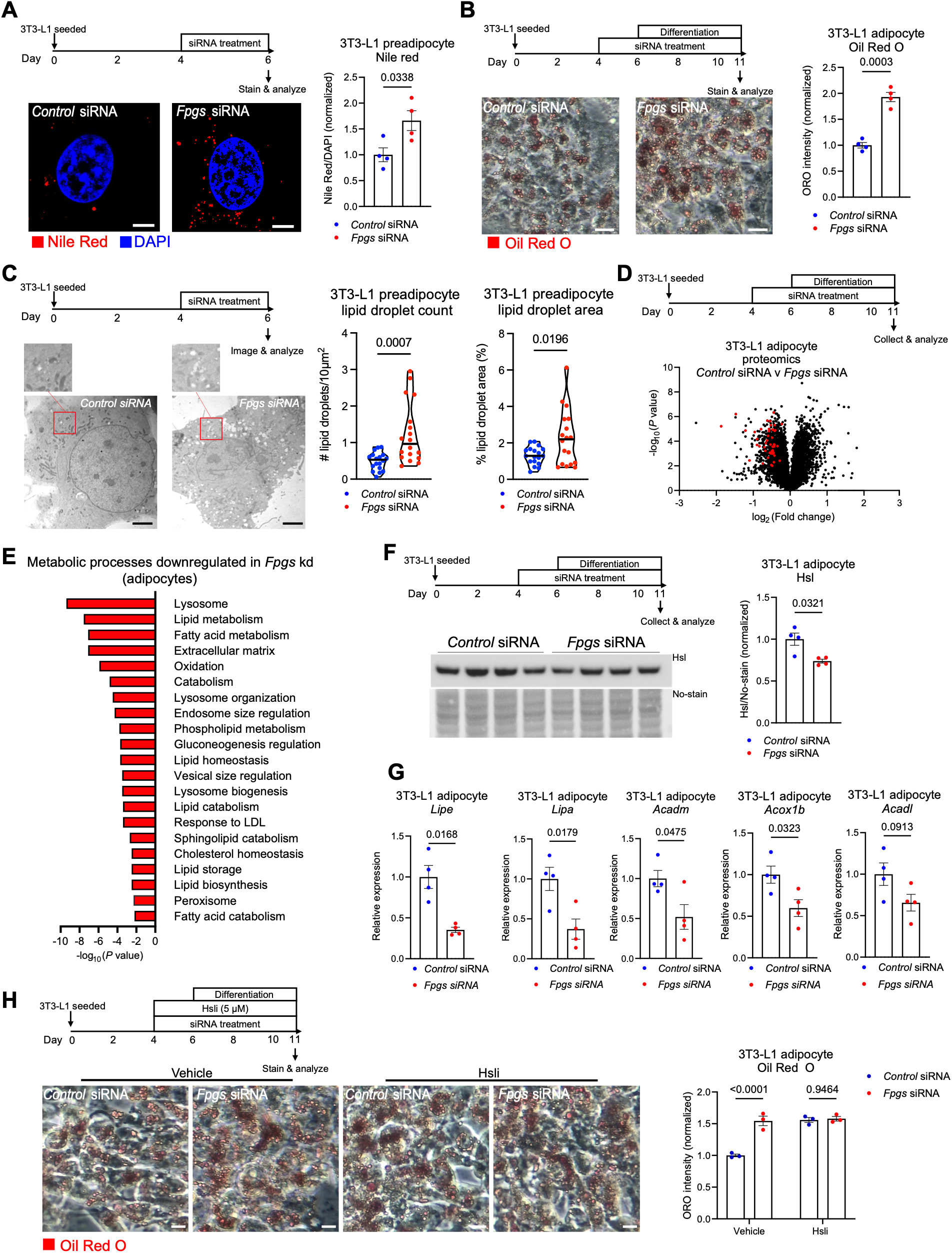
Folic acid polyglutamylation suppresses lipid accumulation via promoting lipid catabolism. (A) Representative images of control and *Fpgs* kd 3T3-L1 preadipocytes stained with Nile Red and DAPI, and quantification (*n* = 4, Welch’s t test. Scale bar = 5 µm). (B) Representative images of control and *Fpgs* kd 3T3-L1 adipocytes stained with Oil Red O, and quantification (*n* = 4, Welch’s t test. Scale bar = 10 µm). (C) Transmission electron microscopy images of control and *Fpgs* kd 3T3-L1 preadipocytes, and quantification (*n* = 18, Mann-Whitney test for droplet count, Welch’s t test for droplet area. Scale bar = 4 µm). (D) Volcano plot displaying abundances of differentially expressed proteins in *Control* and *Fpgs* siRNA treated 3T3-L1 adipocytes. Significantly shifted lipid catabolism-related genes are shown in red. (E) Gene ontology (GO) biological process enrichment analyses of downregulated proteins in *Fpgs* kd (fold change > 1.25, *P* < 0.05). (F) Western blot for hormone sensitive lipase (Hsl) in *Control* and *Fpgs* siRNA treated 3T3-L1 adipocytes (*n* = 4, Welch’s t test). (G) Lipid catabolism genes (*Lipe*, *Lipa*, *Acadm*, *Acox1b, Acadl*) transcript levels in control and *Fpgs* kd cells (*n* = 4, Welch’s t test). (H) Representative images of control and *Fpgs* kd 3T3-L1 adipocytes treated with Hsli (Hi 76-0079), stained with Oil Red O (*n* = 3, Welch’s t test. Scale bar = 10 µm). Bar graphs are represented as mean ± SEM. Violin plots depict data distribution, with the central line indicating the median. All data points represent biological replicates. Exact *P* values are shown in the graphs.

To test whether this increase in lipid accumulation reflects reduced flux through the folate one-carbon cycle rather than the specific loss of FA polyglutamylation, we genetically depleted dihydrofolate reductase (Dhfr), a gateway enzyme that converts FA to downstream reduced folate vitamers including 5-m-THF^21^ (Figure S2C). In contrast to *Fpgs* kd, *Dhfr* kd did not increase lipid accumulation (Figure S2D). Pharmacological inhibition of Dhfr with 100 nM methotrexate^22^ similarly had no effect on lipid levels (Figure S2C and S2E). Together with our metabolomics analysis (Figure 1), these findings suggest that FA polyglutamylation, rather than the canonical downstream folate one-carbon flux, suppresses lipid accumulation in adipocytes.

To investigate the molecular mechanisms driving increased intracellular lipid accumulation following *Fpgs* depletion, we performed unbiased bulk proteomic profiling on control and *Fpgs* kd 3T3-L1 preadipocytes and adipocytes. Global proteomic profiling revealed that Fpgs depletion reshapes adipocyte physiology, upregulating biosynthetic and immune signaling pathways alongside a marked suppression of lipid metabolism in both preadipocytes and adipocytes (Figure S3A-D and Figure 2D, E). Strikingly, *Fpgs* deficiency decreased the abundance of numerous proteins involved in lipid catabolism in both preadipocytes and adipocytes (Figure 2D and Figure S3B). Gene ontology analysis confirmed a selective downregulation of pathways associated with lipid catabolic processes in *Fpgs* kd cells^23^ (Figure 2E and Figure S3C). Consistent with these global proteomics shifts, protein levels of hormone-sensitive lipase (Hsl), a major orchestrator of lipid mobilization^24^, were markedly reduced in *Fpgs* kd cells (Figure 2F). Parallel qPCR analyses revealed a corresponding decrease in the transcripts of key lipid catabolic genes, including the lipase-encoding *Lipe* and *Lipa*, as well as the *β*-oxidation enzymes *Acadm*, *Acadl*, and *Acox1b* (Figure 2G)^25^. Together, these data indicate that Fpgs deficiency transcriptionally represses the core enzymatic machinery of lipid catabolism. Importantly, adipogenic differentiation capacity, assessed by the expression of the master adipogenic regulator peroxisome proliferator-activated receptor-γ (*Pparg*) and transcription co-activators CCAAT/enhancer-binding protein α and β (*Cebpa* and *Cebpb*)^26^, was either unaffected or decreased in *Fpgs* kd cells (Figure S3E and S3F), demonstrating that the observed lipid accumulation does not stem from accelerated adipogenesis. To confirm that compromised lipid breakdown functionally drives this phenotype, we pharmacologically blocked lipid breakdown using the selective Hsl inhibitor Hi 76-0079^27^. This intervention reversed the phenotype, abolishing the difference in lipid accumulation between control and *Fpgs*-deficient cells (Figure 2H). Collectively, these findings demonstrate that FA polyglutamylation regulates lipid accumulation and promotes lipid catabolism in adipocytes.

### Adipose *Fpgs* deficiency suppresses lipid catabolism and induces adipocyte hypertrophy in mice

Next, we investigated whether inhibition of folate polyglutamylation in adipose tissue alters adipose lipid metabolism and whole-body physiology *in vivo*. Using an adipose tissue-targeting Adeno-Associated Virus 8 (AAV8)^28^ encoding eGFP and an shRNA against *Fpgs*, we knocked down the *Fpgs* gene in adipose tissues of chow-fed mice (Figure S4A). Successful transduction of AAV8 was confirmed by an eGFP signal in adipose tissues in a pilot experiment (Figure S4B) and the knockdown efficiency was evaluated by qPCR (Figure S4C). The highest eGFP intensity and a 50% reduction in *Fpgs* expression was observed in iWAT, with subtle shifts in other adipose depots (Figure S4B and S4C). To measure metabolic shifts induced by *Fpgs* kd, mice were monitored for 3.5 months post-AAV8 injection (Figure 3A). By fourteen weeks, the *Fpgs* kd mice exhibited significantly higher body weight gain compared to the control mice (Figure 3B, Figure S4D and S4E), which was not accompanied by an increase in food consumption (Figure S4F). Longitudinal body composition analysis by EchoMRI revealed that this weight increase was accompanied by a significant increase in fat mass (Figure 3C and Figure S4G) as well as a concurrent, significant increase in lean mass in *Fpgs* kd mice (Figure 3D and Figure S4H). Although absolute lean mass expanded, the overall change in body composition was predominantly driven by the increase in fat mass (Figure 3E). Correspondingly, iWAT tissue weight was significantly increased in *Fpgs* kd animals compared to controls (Figure 3F), whereas liver weights remained unchanged (Figure S4I). Collectively these data suggest that suppression of folate polyglutamylation in adipose tissue leads to increased body weight primarily due to adipose tissue expansion.

**Figure 3.**
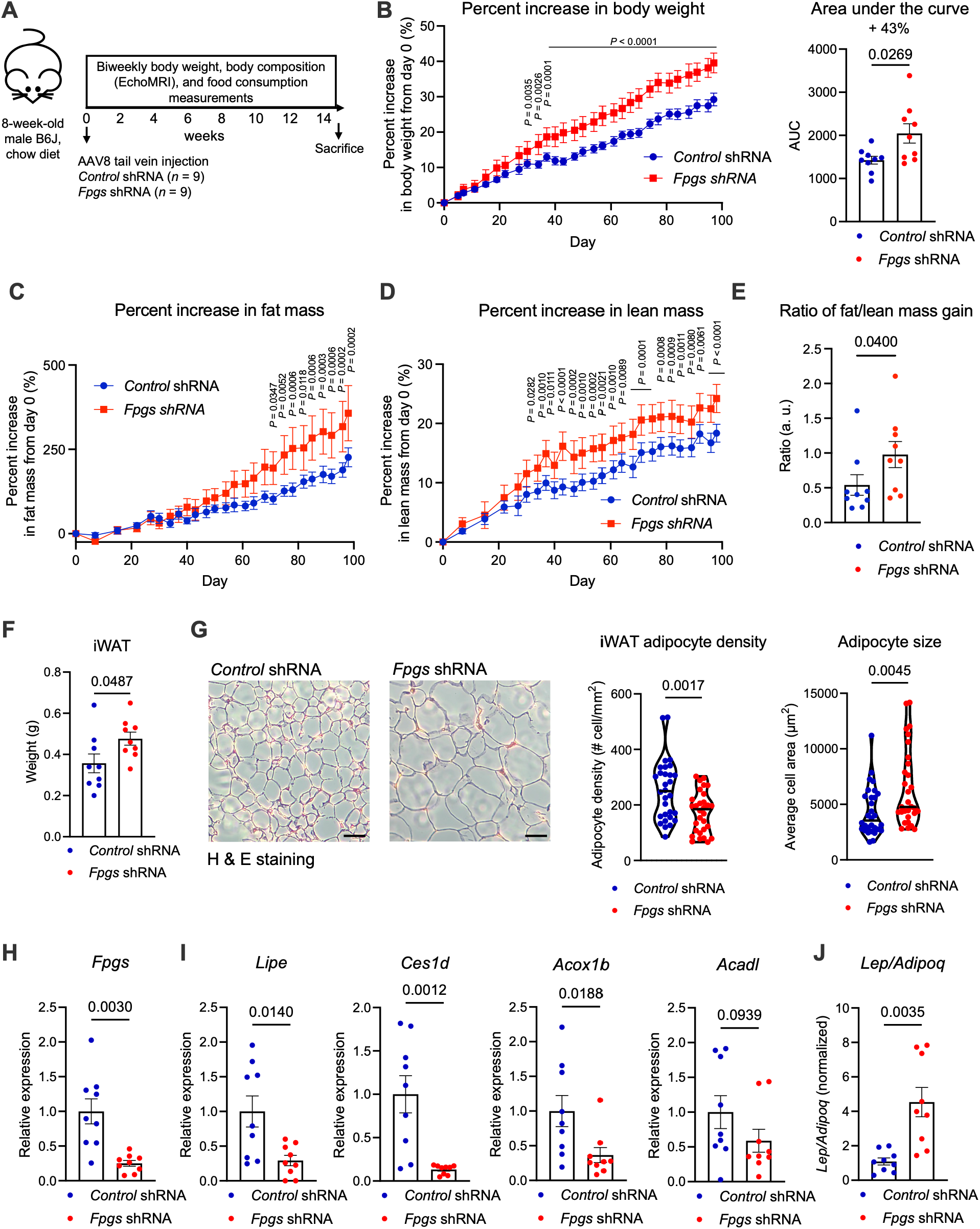
Adipose *Fpgs* knockdown suppresses lipid catabolism and induces adipocyte hypertrophy in mice. (A) Schematic of mouse experiment and timelines. (B) Percent increase in body weight over time and area under the curve (AUC) quantification of *Control* and *Fpgs* kd mice (*n* = 9, Two way ANOVA - multiple comparisons for time course, Welch’s t test for AUC). (C) Percent increase in fat mass over time in *Control* and *Fpgs* kd mice (*n* = 9, Two way ANOVA-multiple comparisons). (D) Percent increase in lean mass over time in *Control* and *Fpgs* kd mice (*n* = 9, Two way ANOVA-multiple comparisons). (E) Ratio of fat mass gain to lean mass gain in *Control* and *Fpgs* kd mice (*n* = 9, Mann-Whiteny test) (F) iWAT weight of *Control* and *Fpgs* kd mice (*n* = 9, Welch’s t test). (G) Representative H&E staining images of iWAT from *Control* and *Fpgs* kd mice, and quantifications (*n* = 30, Welch’s t test for adipocyte density, Mann-Whitney test for adipocyte size. Scale bar = 100 µm). (H, I) Gene transcript levels in control and *Fpgs* kd mice iWAT (*n* = 9, Welch’s t test for *Fpgs*, *Lipe*, Mann-Whitney test for *Acox1b*, *Ces1d*). (J) Normalized *Lep/Adipoq* ratio in iWAT of control and *Fpgs* kd mice (*n* = 9, Welch’s t test). Bar graphs are represented as mean ± SEM. All data points represent biological replicates. Time course plots are represented as mean ± SEM. Violin plots depict data distribution, with the central line indicating the median. Data points represent H&E images taken in a blinded manner. Exact *P* values are shown in the graphs. Data not marked with exact *P* values are not significant (*P* > 0.05).

To investigate the cellular consequences of increased adiposity, we examined adipose tissue histology. iWAT sections were stained with hematoxylin and eosin (H&E) to quantify adipocyte surface area. Strikingly, adipocytes from *Fpgs* kd mice were significantly more hypertrophic than those from control mice (Figure 3G). Consistent with our *in vitro* and pilot *in vivo* findings, *Fpgs* kd led to a ∼75% decrease in *Fpgs* mRNA levels in iWAT (Figure 3H), which was accompanied by a reduction in the expression of lipid catabolism genes (*Lipe*, *Ces1d*, *Acox1b*, *Acadl*) without altering expression of adipogenic markers fatty acid synthase (*Fasn*) and *Pparg* (Figure 3I and Figure S4J). Furthermore, the iWAT leptin-to-adiponectin transcript ratio (L:A), an indicator of adipose tissue dysfunction^29^, was significantly higher in *Fpgs* kd mice (Figure 3J and Figure S4K). Together, these findings demonstrate that folate polyglutamylation is required to maintain adipose lipid catabolism *in vivo*, and that loss of this pathway promotes adipocyte hypertrophy, coordinated adipose tissue expansion, and body weight gain even in the absence of a dietary challenge.

### FA monoglutamate binds and inhibits mitogen activated protein kinase kinase 5 (Map2k5)

We next sought to elucidate the molecular mechanisms driving the repression of lipid catabolic gene expression in *Fpgs* kd cells. Folate deficiency has previously been linked to altered transcriptional patterns through DNA instability and epigenetic methylation changes^30^. However, we did not observe changes to OCM in WATs in obesity that significantly correlated with body weight (Figure S1D). Further, metabolic alterations typically require days or weeks to impair the epigenome or structural DNA integrity^31,32^. This extended timeline is incompatible with the rapid, 2-day post-knockdown phenotype we observed (Figure 2A), pointing to a direct, alternative pathway regulating gene expression.

Given that *Fpgs* kd prevents folate polyglutamylation, we reasoned that folates differing in their polyglutamylation states might differently interact with signaling enzymes to modulate gene transcription. To profile such interactions, we screened for binding partners of FA-monoglutamate (FA-1G), -diglutamate (FA-2G), -tetraglutamate (FA-4G), and -hexaglutamate (FA-6G) in 3T3-L1 preadipocytes using target engagement proteomics (TEP) (Figure S5A)^33^. Known folate polyglutamate-dependent enzymes such as Dhfr^34^ and Atic^35^, displayed a progressive increase in thermal stability with extending glutamyl chain lengths, thus validating our TEP platform (Figure 4A). Conversely, at the opposite spectrum of the interaction profile, we identified mitogen-activated protein kinase kinase 5 (Map2k5) as a top candidate that exhibited a successive decrease in binding stability as the folic acid glutamyl chain length increased. A ligand binding assay using recombinant purified human MAP2K5 with LC-MS analysis confirmed the physical interaction between FA and MAP2K5 (Figure 4B). Orthogonal limited proteolysis revealed that incubation with FA increased the protease susceptibility of MAP2K5, consistent with an FA-induced conformational change upon binding (Figure 4C).

**Figure 4.**
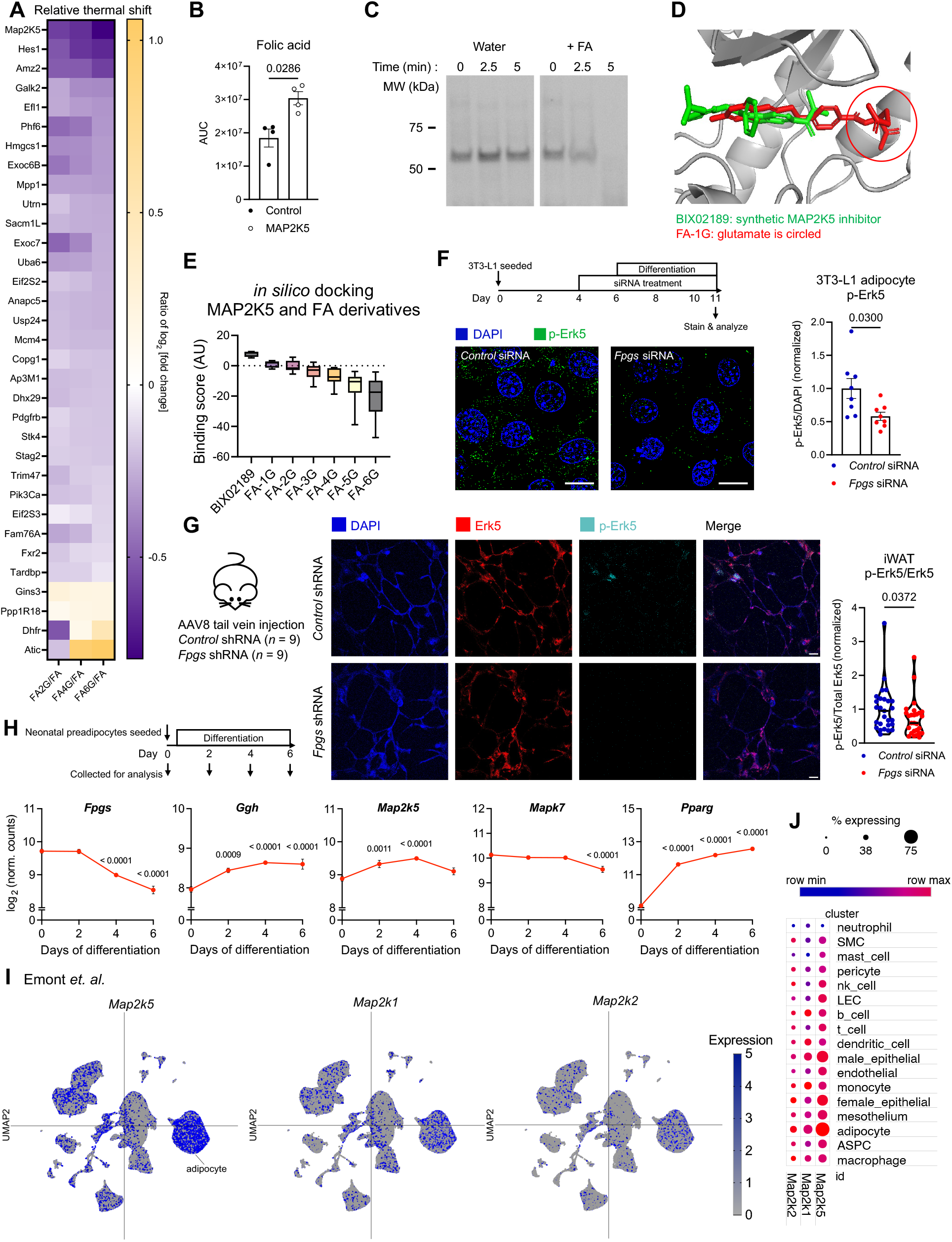
Folic acid monoglutamate binds and inhibits mitogen activated protein kinase kinase 5 (Map2k5). (A) Heat map displaying relative thermal stability shift of proteins (in rows) when they are bound to polyglutamylated folic acids (FA-2/4/6G, in columns) compared to when bound to monoglutamylated folic acid (FA-1G). (B) LC-MS quantification of FA extracted from no-protein control and purified MAP2K5 bound beads (*n* = 4, Mann-Whitney test). (C) Limited proteolysis of purified MAP2K5 with control or FA for the indicated time points. (D) Docking simulation of BIX02189 (green) and FA-1G (red) bound to MAP2K5. Glutamate residue in FA-1G is circled. (E) Calculated binding affinity of BIX02189 and FA with varying polyglutamyl chain length. Binding score = -SMINA affinity score. (F) Immunocytochemistry displaying phosphorylated Erk5 (green) and nucleus (blue) in control and *Fpgs* kd 3T3-L1 adipocytes (*n* = 8, Welch’s t test. Scale bar = 10 µm). (G) Representative images of mouse iWAT stained with antibodies against Erk5, phosphorylated Erk5 and DAPI, with quantification for Erk5 phosphorylation rate (*n* = 27, Mann-Whitney test. Scale bar = 10 µm). (H) Gene expression levels along white adipocyte differentiation timeline *ex vivo* (*n* = 3, one-way ANOVA - multiple comparisons test). (I) snRNA seq data in whole mouse adipose tissue from Emont et. al. for indicated transcripts. (J) Dot plot showing expression level of indicated transcripts in whole mouse adipose tissue. All bar graphs are represented as mean ± SEM, data points represent biological replicates. Floating bars represent the top 10 docking simulation scores. Exact *P* values are shown in the graphs.

To investigate the structural basis of this interaction, we performed molecular docking simulations^36^ between the Map2k5 enzyme and FA species ranging from monoglutamylated to hexaglutamylated forms (Figure 4D). These simulations revealed that all FA species occupy the same binding pocket within Map2k5, but exhibit a progressive decrease in binding affinity as the glutamyl chain elongates, driven primarily by steric hindrance (Figure 4D and 4E). Interestingly, BIX02189, a known selective synthetic inhibitor of Map2k5 kinase activity^37^, was also predicted to dock within the same binding pocket, suggesting that FA-1G may function as an endogenous Map2k5 inhibitor. In the canonical MAP2K5-ERK5 signaling pathway, MAP2K5 phosphorylates and activates ERK5 (MAPK7)^38^. To test whether the attenuation of folate polyglutamylation inhibits Map2k5 signaling in cells, we probed for phosphorylated Erk5 (p-Erk5) and total Erk5 levels in control and *Fpgs* kd 3T3-L1 adipocytes by immunocytochemistry and western blot analyses. p-Erk5 levels were significantly downregulated in *Fpgs* kd cells, while total Erk5 levels remained unchanged (Figure 4F, Figure S5B and S5C). Importantly, this signaling suppression translated *in vivo*, where *Fpgs* kd iWAT displayed a concomitant reduction in p-Erk5 levels. (Figure 3A and 4G). Together, these findings demonstrate that suppression of folate polyglutamylation leads to the accumulation of monoglutamated FA, which targets Map2k5 to inhibit Erk5 signaling both *in vitro* and *in vivo*.

### Map2k5 activity promotes lipolytic gene expression to prevent adipocyte hypertrophy

Because the relationship between folate polyglutamylation and Map2k5 signaling remains uncharacterized in adipocyte lipid metabolism, we measured transcript levels of key components of this signaling axis in differentiating white adipocytes *ex vivo^39^*. *Fpgs* expression decreased, whereas *Ggh* transcript levels increased during differentiation, consistent with reduced folate polyglutamylation as adipocytes accumulate lipids and shift toward lipid storage over catabolism. *Map2k5* and *Erk5* (*Mapk7*) were highly expressed in preadipocytes, at levels comparable to *Pparg*, and remained elevated throughout differentiation (Figure 4H), consistent with our observations across the 3T3-L1 differentiation timeline (Figure 2A-C). Next, we interrogated a publicly available single-nuclear (snRNA-seq) transcriptomic dataset^40^ to determine the physiological relevance and expression profile during differentiation of Map2k5 compared to other canonically studied Map2k proteins in adipocytes. Analysis of *in vivo* snRNA-seq data from mouse white adipose tissue confirmed high tissue-wide *Map2k5* expression, with the highest abundance localized to mature adipocytes (Figure 4I and 4J). Notably, *Map2k5* transcript levels substantially exceeded those of *Map2k1* and *Map2k2*, which encode for the canonical MEK kinases Mek1 and Mek2 respectively that operate in the classical adipocyte insulin signaling pathway. Together, these data establish Map2k5 as a highly expressed, cell-type-enriched kinase in adipose tissues, identifying it as a downstream target of folate polyglutamylation.

We next evaluated the functional requirement for Map2k5 in adipocyte lipid metabolism. Given that FA-1G inhibits Map2k5, we reasoned that suppressing Map2k5 signaling alone would be sufficient to drive intracellular lipid accumulation. Pharmacological inhibition of Map2k5 with a selective synthetic inhibitor BIX02189 significantly increased lipid content in both preadipocytes (Figure 5A) and differentiated adipocytes (Figure S6A). Genetic depletion of *Map2k5* via siRNA transfection similarly triggered marked lipid accumulation in 3T3-L1 adipocytes (Figure 5B). Mechanistically, both pharmacological inhibition (Figure 5C), and genetic depletion of *Map2k5* downregulated the expression of key lipid catabolic genes without affecting markers of adipogenesis (Figure 5D and Figure S6B). Together, these data demonstrate that Map2k5 signaling promotes lipid catabolism and limits pathological lipid accumulation in adipocytes.

**Figure 5.**
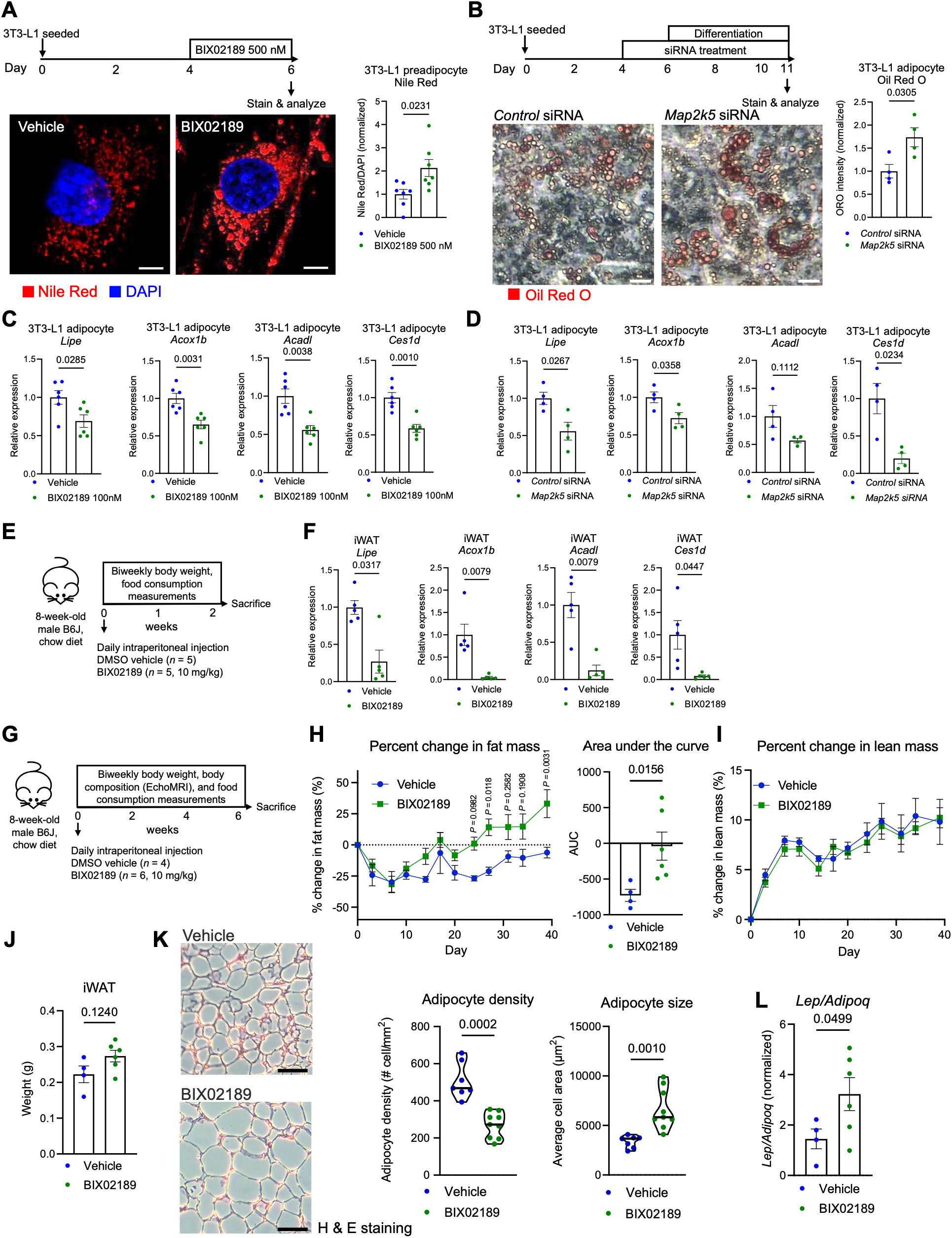
Map2k5 activity promotes lipolytic gene expression to prevent adipocyte hypertrophy. (A) Representative images of vehicle and BIX02189 treated 3T3-L1 preadipocytes stained with Nile Red and DAPI, and quantification (*n* = 7, Welch’s t test. Scale bar = 5 μm). (B) Representative images of control and *Map2k5* kd 3T3-L1 adipocytes stained with Oil Red O (*n* = 4, Welch’s t test. Scale bar = 10 μm). (C) Lipid catabolism gene transcript levels in vehicle and BIX02189 treated cells (*n* = 6, Welch’s t test). (D) Lipid catabolism gene transcript levels in control and *Map2k5* kd 3T3-L1 adipocytes (*n* = 4, Welch’s t test). (E) Schematic of acute BIX02189 treatment mouse experiment and timeline. (F) Lipid catabolism gene transcript levels in iWAT of vehicle and BIX02189 treated mice (*n* = 5, Welch’s t test for *Ces1d*, Mann-Whitney test for all other genes). (G) Schematic of chronic BIX02189 treatment mouse experiment and timeline. (H) Percent change in fat mass over time of vehicle and BIX02189 treated mice and area under the curve quantification (*n* = 4 for vehicle, 6 for BIX02189, Two way ANOVA - multiple comparisons for time course, Welch’s t test for AUC). (I) Percent change in lean mass over time of vehicle and BIX02189 treated mice (*n* = 4 for vehicle, 6 for BIX02189, Two way ANOVA - multiple comparisons). (J) iWAT weight (*n* = 4 for vehicle, 6 for BIX02189, Welch’s t test). (K) Representative H&E staining images of iWAT from vehicle and BIX02189 treated mice and quantifications (*n* = 7 for vehicle, 9 for BIX02189, Welch’s t test, scale bar is 100 μm). (L) Normalized *Lep/Adipoq* ratio in iWAT of vehicle and BIX02189 treated mice (*n* = 4 for vehicle, 6 for BIX02189, Welch’s t test). Bar graphs are represented as mean ± SEM. All data points represent biological replicates. Time course plots are represented as mean ± SEM. Time course plots are represented as mean ± SEM. Violin plots depict data distribution, with the central line indicating the median. Data points represent H&E images taken in a blinded manner. Exact *P* values are shown in the graphs. Data not marked with exact *P* values are not significant (*P* > 0.05).

Next, we evaluated whether Map2k5 activity is required for adipocyte lipid catabolism *in vivo*. Daily intraperitoneal (i.p.) administration of BIX02189 for two weeks (Figure 5E) significantly downregulated lipid catabolic gene expression in iWAT (Figure 5F), underscoring the physiological relevance of Map2k5 signaling *in vivo*. BIX02189-treated mice exhibited a subtle upward trend in body weight (Figure S6C-E), without alterations in food intake (Figure S6F). To assess the physiological consequences of sustained Map2k5 suppression, we extended BIX02189 administration to 6 weeks (Figure 5G). Similarly to the acute experiment, chronic Map2k5 inhibition maintained a trend toward increased body weight without affecting caloric consumption (Figure S6G-J). Body composition analysis revealed a significant increase in total fat mass (Figure 5H and Figure S6K) without a change in lean mass (Figure 5I and Figure S6L), suggesting that Map2k5 inhibition can increase adiposity. Although total iWAT mass weight increase following BIX02189 treatment did not reach statistical significance (Figure 5J), histological evaluation of tissue sections by H&E revealed marked adipocyte hypertrophy (Figure 5K), which was reflected by a higher L:A ratio (Figure 5L and Figure S6M). Together, these findings demonstrate that Map2k5 is a critical regulator of adipocyte lipid homeostasis both *in vitro* and *in vivo*.

### FPGS and MAP2K5 are negatively correlated with obesity markers in humans

To test whether folate polyglutamylation regulates lipid metabolism in humans, we knocked down *FPGS* in an immortalized human adipocyte cell line. *FPGS*-kd adipocytes displayed significantly increased lipid accumulation compared to control cells (Figure 6A), demonstrating a conserved role for *FPGS* in preventing lipid accumulation across species.

**Figure 6.**
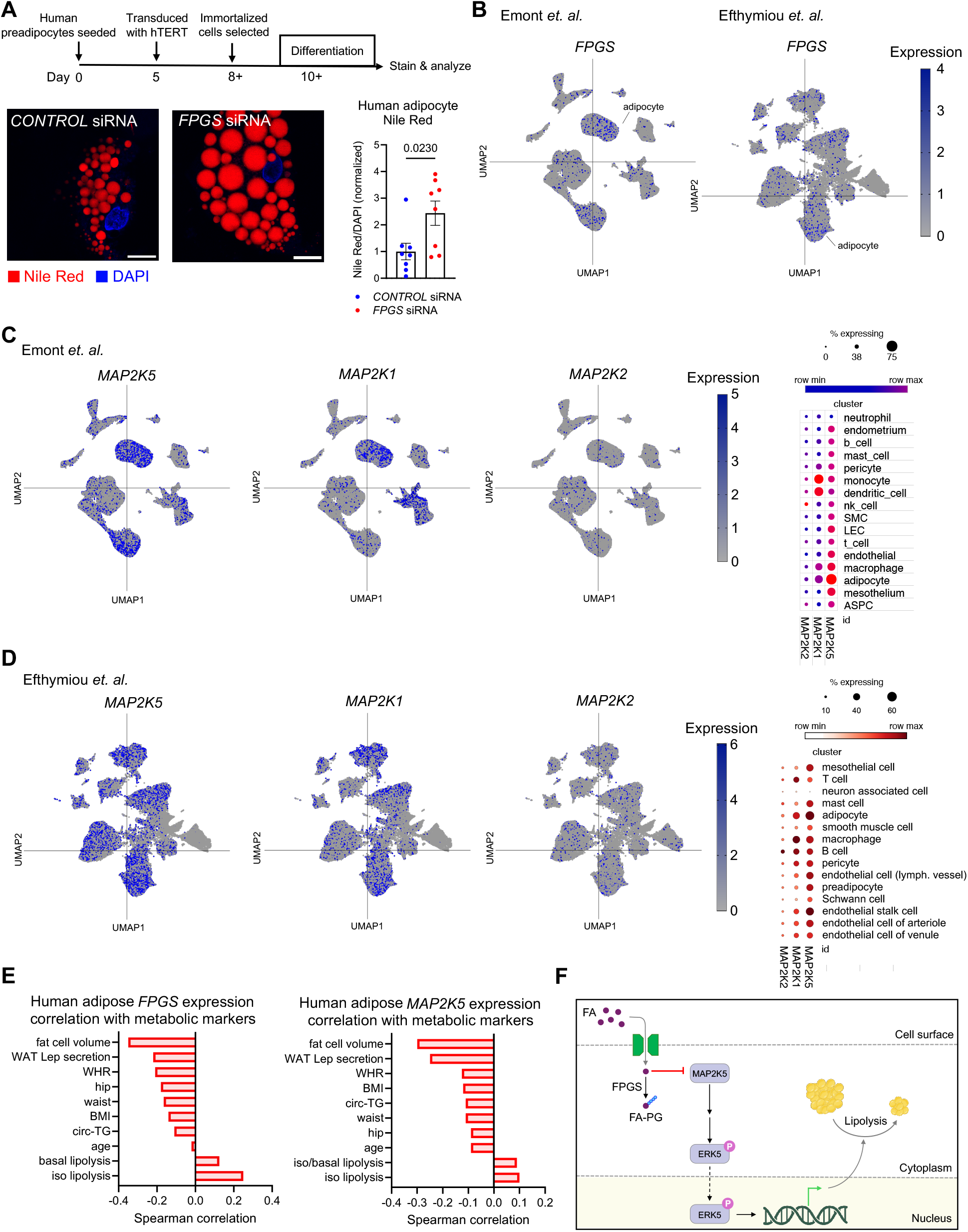
*FPGS* and *MAP2K5* are negatively correlated with obesity markers in humans. (A) Representative images of control and *Fpgs* kd immortalized human adipocytes stained with Nile Red and DAPI and quantification (*n* = 4, Welch’s t test, Scale bar is 10 μm). (B-D) UMAPs displaying *FPGS*, *MAP2K1, MAP2K2,* and *MAP2K5* expression in human adipose tissue. (E) Correlation between *FPGS* and *MAP2K5* transcript levels in human adipose tissues and clinical metabolic health markers. (F) Model. All bar graphs are represented as mean ± SEM, data points represent biological replicates. Exact *P* values are shown in the graphs.

To establish the translational relevance of these findings, we queried publicly available human single-cell transcriptomic datasets and clinical cohorts. Analysis of snRNA-seq profiles from visceral and subcutaneous adipose tissues from 13 individuals revealed that *FPGS* and *MAP2K5* are expressed across the adipose tissue subpopulations including high expression in adipocytes^40^ (Figure 6B-D). This distribution was corroborated in another data set from 22 human subjects’ subcutaneous and intraabdominal adipose tissues^41^ (Figure 6B-D). Consistent with our murine snRNA-seq findings (Figure 4I), *MAP2K5* was the predominant MEK isoform expressed in human adipose tissue, surpassing *MAP2K1* and *MAP2K2*. Finally, interrogation of the Adipose Tissue Data Portal^42^ demonstrated that *FPGS* and *MAP2K5* transcript levels inversely correlate with clinical indices of obesity, while positively correlating with *ex vivo* adipocyte lipolytic capacity (Figure 6E). Together, these clinical and human cell-based findings establish the *FPGS-MAP2K5* axis (Figure 6F) as a conserved, anti-obesogenic metabolic checkpoint with high translational relevance for human metabolic health.

## DISCUSSION

This study identifies a previously unrecognized signaling axis linking micronutrient processing directly to adipocyte function. While one-carbon units supplied by folates are vital for metabolic maintenance, our findings uncover a non-canonical, signaling-centric role for vitamin B9 where intracellular folate dynamics directly coordinate kinase signaling cascades in adipose tissue. By demonstrating that FPGS-mediated folate polyglutamylation acts as an endogenous chemical switch to regulate MAP2K5 activity and downstream lipid handling independently of canonical OCM flux, our findings redefine folate from a passive metabolic substrate to an active, non-canonical regulator of adipocyte homeostasis across species.

Historically, clinical and preclinical assessments of folate status have relied almost exclusively on circulating blood levels^43^. However, systemic metrics fail to reflect local tissue dynamics, as evidenced by the marked heterogeneity in folate vitamer distribution across mouse tissues^18^. This study further highlights the divergence between systemic and adipose depot-specific folate pools. While systemic folate levels exhibit only modest shifts in obesity and weight loss, resident adipose tissue folates suffer a substantial and selective shift in mice and humans in response to weight changes. The obesity-associated depletion of folate demonstrates that adipose tissue operates under distinct homeostatic controls, revealing local folate deficiency as a central feature of tissue dysfunction.

Folate polyglutamylation by FPGS is an evolutionarily conserved biochemical process operating across bacteria, plants, and animals. It is essential for cellular folate retention and enhancing binding affinity with OCM enzymes. Complete disruption of Fpgs, thus, causes embryonic lethality in animals^44^, and severe proliferative arrest in cultured cells^45^, underscoring its foundational biochemical role. Considering the critical role of OCM in methylation reactions and nucleotide synthesis, it has long been linked to obesity and cardiometabolic diseases^9^. Previous studies have shown that folate availability can modulate the methylation status of hypertrophy-related genes in adipocytes^17^. Notably, while we observed no immediate shifts in lipid storage following acute Dhfr silencing or methotrexate treatment, chronic methotrexate exposure has been reported to drive lipid accumulation^46^. These previous findings underscore how chronic folate deficiency may govern long term adipocyte metabolism via epigenetic reprogramming. Here, we uncover a distinct, parallel mechanism wherein monoglutamylated folic acid directly binds and regulates MAP2K5 kinase activity to govern lipid handling independently of OCM. How FPGS-mediated folate polyglutamylation intersects with downstream OCM dynamics and genome-wide methylation in adipocytes remains an important avenue for future investigation.

While MAP2K5 has previously been linked to PPARG-mediated adipogenesis^47^, whether it directly governs mature adipocyte lipid handling remained unclear. Because alterations in adipocyte differentiation can influence intracellular lipid accumulation, we sought to distinguish the effects of MAP2K5 on adipogenesis from its effects on lipid metabolism. We therefore maintained PPARG agonist-mediated adipogenic differentiation under standardized conditions while manipulating MAP2K5 signaling, allowing us to assess its effects on lipid metabolism independently of differences in adipocyte differentiation. Under these standardized conditions, both genetic and pharmacological suppression of MAP2K5 increased intracellular lipid accumulation. These data demonstrate that MAP2K5 actively restrains lipid storage independently of PPARG-driven differentiation, establishing a distinct cell-autonomous role for the FPGS-MAP2K5 axis in basal adipocyte lipid catabolism. Given that *MAP2K5* is highly expressed throughout the adipocyte differentiation, further research into the interplay between MAP2K5 and PPARG during adipogenesis and in mature adipocyte metabolism is warranted.

As the direct upstream kinase of ERK5, MAP2K5 has primarily been studied in vascular development and oncology^48^, although human genetic variants have linked MAP2K5 loci to metabolic disease^49^. Prior work has demonstrated that adipose-specific *Mapk7* KO mice develop weight gain under both standard chow and high-fat diet regimens^38^, corroborating our work and reinforcing the importance of the MAP2K5-ERK5 cascade in adipocyte function. The upstream triggers initiating this signaling pathway in adipose tissue are yet to be determined. While mechanosensory receptors trigger MAP2K5–ERK5 signaling in endothelial cells^37^, whether these receptors are expressed in adipocytes is unknown. Both *Fpgs* kd and BIX02189-treated mice exhibited increased adiposity on standard chow without changes in total caloric intake or macronutrient composition. These findings position the FPGS-MAP2K5 axis as an essential, cell-autonomous checkpoint for maintaining basal lipid homeostasis. Coupled with the inverse correlation between MAP2K5 expression and clinical markers of obesity in humans, these insights highlight the FPGS-MAP2K5 cascade as a promising target for therapeutic intervention in metabolic disease.

## RESOURCE AVAILABILITY

### Lead contact

Further information and requests for information and resources should be directed to and will be fulfilled by the lead contact, Snehal N. Chaudhari.

### Materials availability

This study did not generate new unique reagents.

### Data and code availability

- Raw proteomics data have been deposited to the ProteomeXchange Consortium with the dataset identifier PXD083247 for bulk proteomics and PXD083292 for target engagement proteomics. Data are publicly available as of the date of publication.
- This paper analyzes existing, publicly available datasets. The accession numbers for snRNA and scRNA data are listed in the Key Resources Table.
- No custom code was used in this study.
- All other data reported in this paper will be shared by the lead contact upon request.

## ACKNOWLEDGEMENTS

This study was supported by the National Institutes of Health R00 DK128503 (S.N.C), R35GM150899 (A.G.), National Institutes of Health T32 GM152341 (W.S.K.), the Ruth Dickie Endowment from the Madison Chapter of the Graduate Women in Science (J.C.), National Institutes of Health T32 GM135066 (J.W.), National Institutes of Health T32 DK007665 (H.K.), the Wisconsin Alumni Research Foundation (WARF), and the department of Biochemistry at University of Wisconsin-Madison. We are grateful to the participants who donated tissue samples for scientific analyses to the Brigham and Women’s Hospital and the University of Wisconsin Health Organ and Tissue Donation program. We would like to thank the UWCC Experimental Animal Pathology Laboratory for all histology and immunofluorescent staining services, Electron Microscope Facility for TEM images, and K.R. Weiss from the Biochemistry Optical Core at UW-Madison for assistance with fluorescence imaging. We acknowledge the Simcox and Wei labs at UW-Madison for help with equipment and reagents. A special thank you to Drs. Feyza Engin, Judith Simcox, and Alan Attie at UW-Madison for insightful guidance that strengthened this work. Finally, we are grateful to the members of the Chaudhari lab for technical assistance, helpful discussions, and critical reviews.

## AUTHOR CONTRIBUTIONS

Conceptualization and study design was carried out by W.S.K. and S.N.C. W.S.K. performed in all *in vitro* and *in vivo* experiments along with tissue processing and data analysis with the help of R.D., J.C., D.K., J.H. and L.L. Development of the folate metabolomics workflow was performed by J.W., and optimization for adipose tissues and data processing was performed by W.S.K. with the help of L.L. W.S.K. and H.K. performed MAP2K5 bacterial expression, purification, and folate binding assays. J.G.V.V. performed bulk and target engagement proteomics. Western diet feeding and sleeve gastrectomy were performed by D.A.H. Analysis of differentiating adipocyte scRNA was performed by Y.B. and A.G. Immortalized human adipocyte cell line was generated by E.M.P. and B.W.P. W.S.K. and S.N.C. wrote the paper and prepared all figures. All authors edited and contributed to the critical review of the paper.

## SUPPLEMENTAL INFORMATION

Document S1. Figures S1–S6

Table S1. Analytical parameters of folate extraction method.

Table S2. The primer list for qPCR. Related to STAR Methods.

---

**Figure S1.**
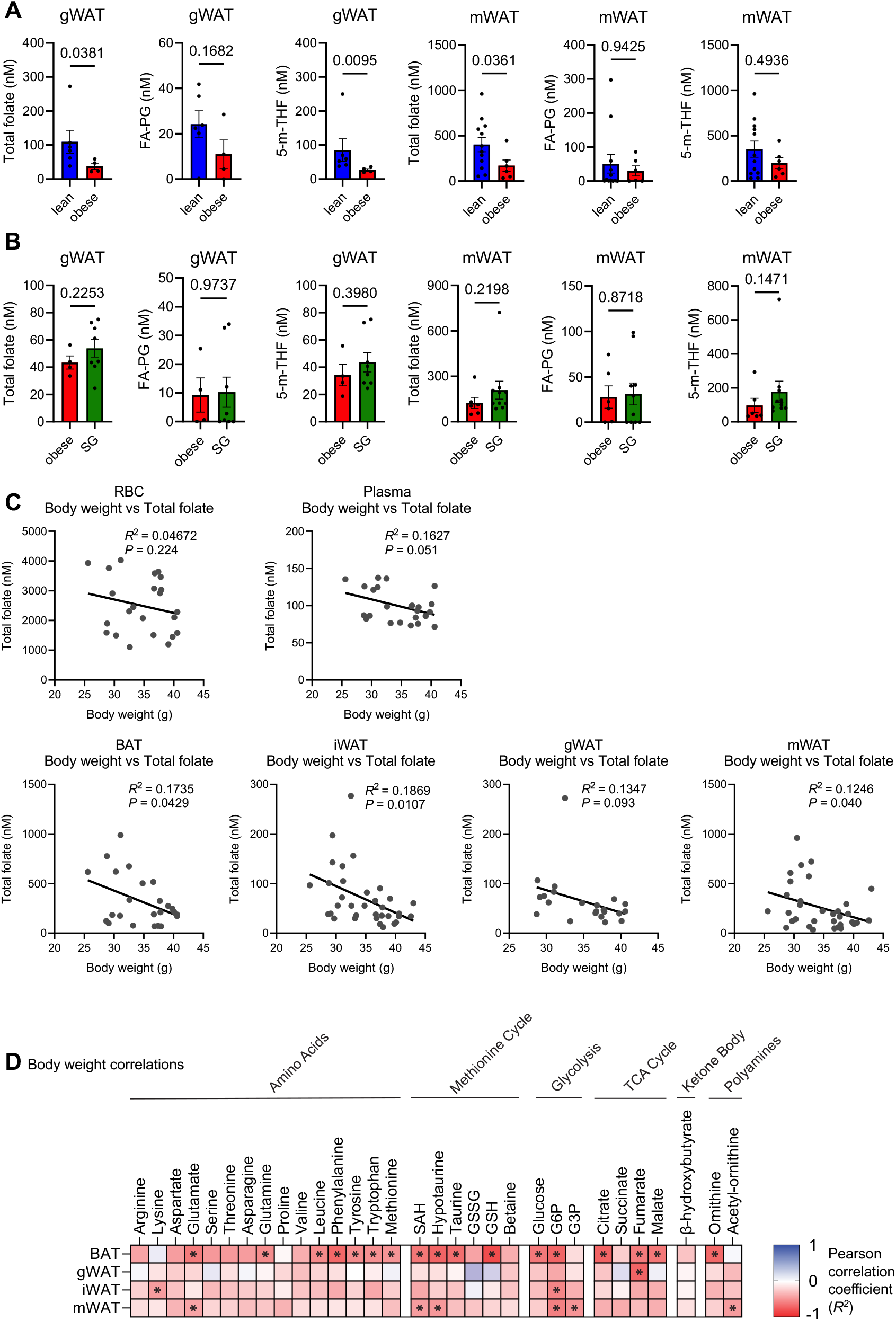
Adipose resident folates are depleted in obesity. Related to Figure 1. (A) Folate levels in gonadal white adipose tissue (gWAT) and mesenteric white AT (mWAT) from lean and obese mice (gWAT: lean *n* = 6, obese *n* = 4; mWAT: lean *n* = 12, obese *n* = 6, Mann-Whitney test for gWAT total folate, 5-m-THF, mWAT FA-PG, 5-m-THF, and Welch’s t test for gWAT FA-PG, mWAT total folate). (B) Folate levels in gWAT and mWAT from obese and SG mice (gWAT: obese *n* = 4, SG *n* = 8; mWAT: obese *n* = 6, SG *n* = 10, Mann-Whitney test for gWAT FA-PG, all mWATs, Welch’s t test for gWAT total folate, 5-m-THF). (C) Correlational analysis between body weight and total folate levels in RBC and plasma (*n* = 24, Pearson correlation, two-sided Student’s t-test) and BAT, iWAT, gWAT, mWAT (*n* = 24, 34, 22, 34, respectively, Pearson correlation, two-sided Student’s t-test). *R*^2^ and *P* values are indicated on the correlation graphs; the curve represents the line of best fit. (D) Heat map of Pearson correlation of indicated metabolites with body weight across adipose tissues. Asterisks indicate a statistically significant correlation (*P* < 0.05) (BAT, iWAT, gWAT, mWAT (*n* = 24, 34, 22, 34, respectively, Pearson correlation, two-sided Student’s t-test). All bar graphs are represented as mean ± SEM, data points represent biological replicates. Exact *P* values are shown in the graphs.

**Figure S2.**
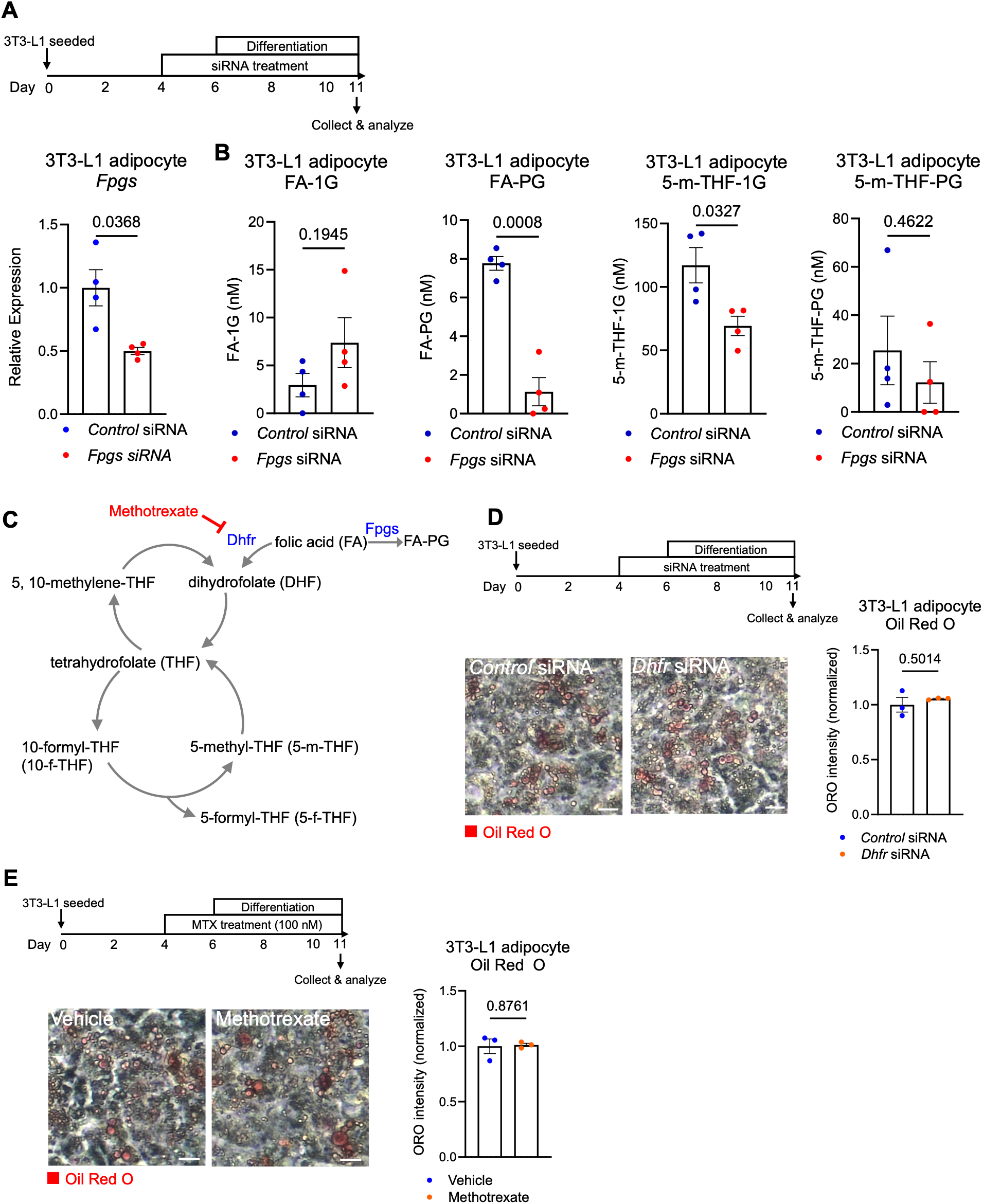
Inhibiting flux into the folate cycle does not induce lipid accumulation. Related to Figure 2. (A) *Fpgs* transcript level in control and *Fpgs* kd 3T3-L1 adipocytes (*n* = 4, Welch’s t test). (B) Levels of folate vitamers in control and *Fpgs* kd 3T3-L1 adipocytes (*n* = 4, Welch’s t test). (C) Folate cycle (D, E) Oil Red O staining levels in 3T3-L1 adipocytes treated with (D) *control* and *Dhfr* siRNA (*n* = 3, Welch’s t test, Scale bar = 10 µm) or (E) vehicle and Methotrexate (100 nM) (*n* = 3, Welch’s t test. Scale bar = 10 µm). All bar graphs are represented as mean ± SEM, data points represent biological replicates. Exact *P* values are shown in the graphs.

**Figure S3.**
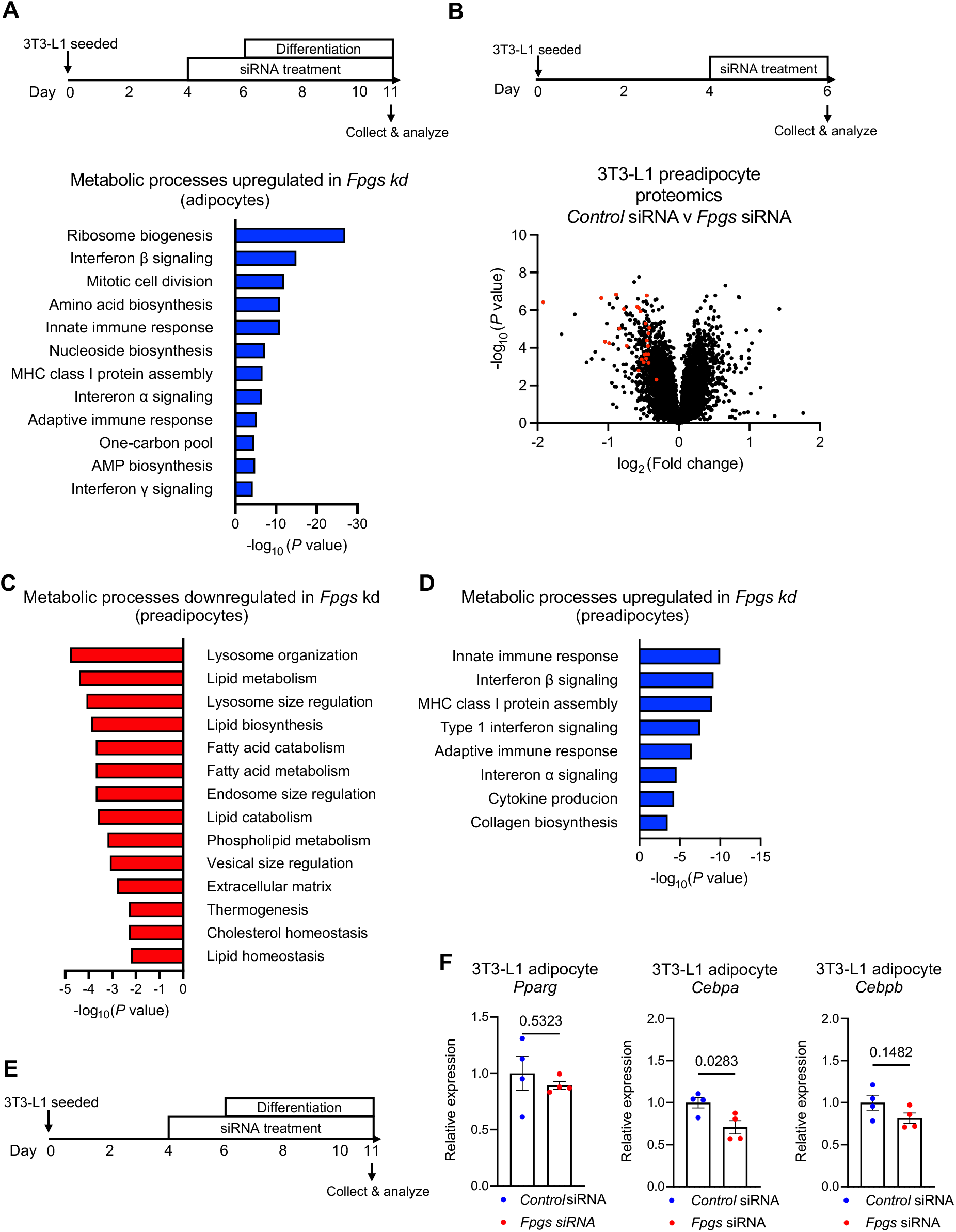
Folic acid polyglutamylation suppresses lipid accumulation via promoting lipid catabolism. Related to Figure 2. (A) Gene ontology (GO) biological process enrichment analyses of upregulated proteins in 3T3-L1 adipocyte *Fpgs* kd cells (fold change > 1.25, *P* < 0.05). (B) Volcano plot displaying abundances of differentially expressed proteins in *Control* and *Fpgs* siRNA treated 3T3-L1 preadipocytes. Significantly shifted lipid catabolism-related genes are shown in red. (C, D) Gene ontology (GO) biological process enrichment analyses of downregulated proteins (C) and upregulated proteins (D) in 3T3-L1 preadipocyte *Fpgs* kd cells (fold change > 1.25, *P* < 0.05). (E) Experimental timeline. (F) Adipogenesis marker gene transcript levels in *Control* and *Fpgs* kd cells (*n* = 4, Welch’s t test). All bar graphs are represented as mean ± SEM, data points represent biological replicates. Exact *P* values are shown in the graphs.

**Figure S4.**
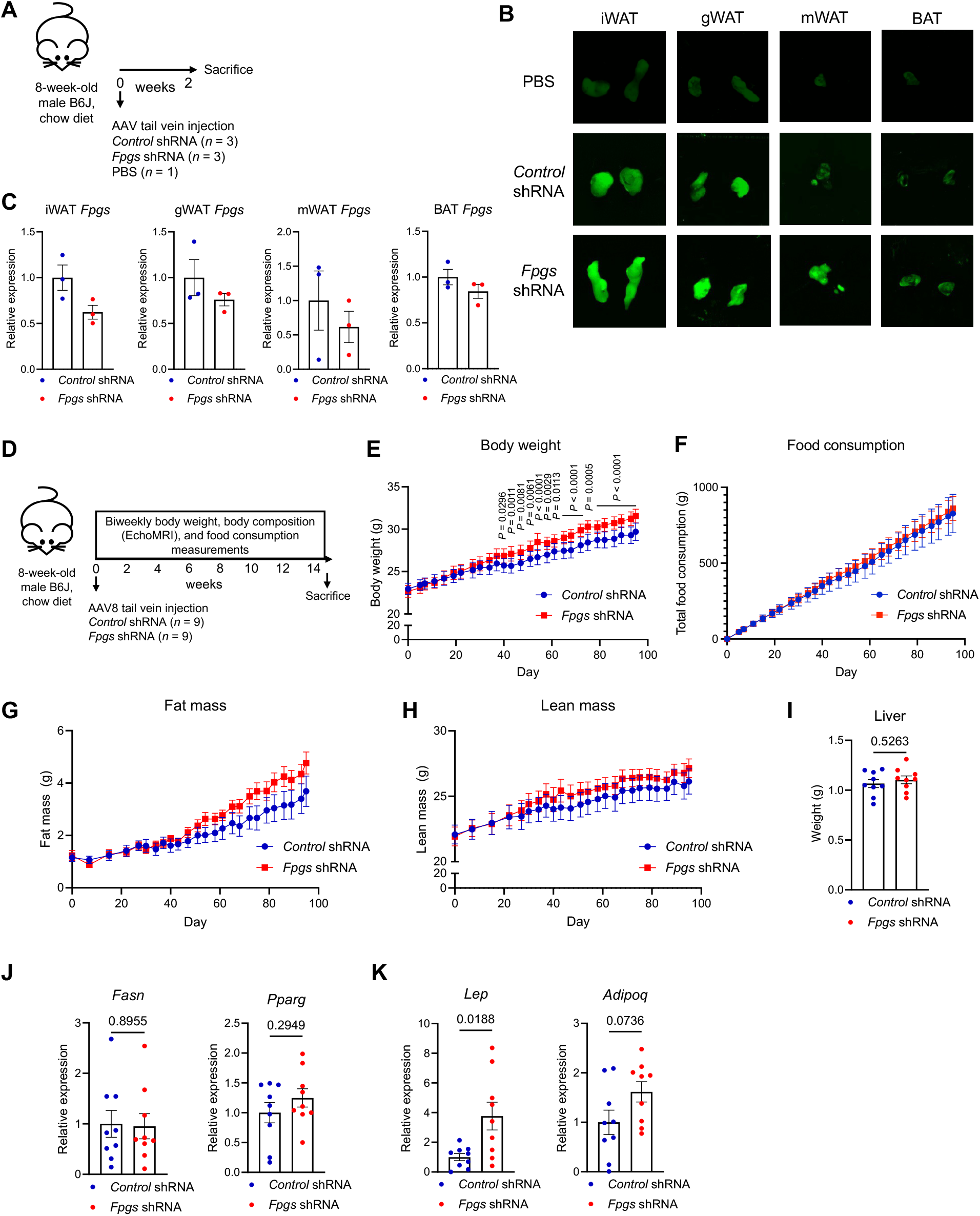
Adipose *Fpgs* knockdown suppresses lipid catabolism and induces adipocyte hypertrophy in mice. Related to Figure 3. (A) Schematic of pilot mouse experiment and timeline. (B) Fluorescence images showing eGFP signals *ex vivo* in iWAT, gWAT, mWAT, and BAT of mice injected with PBS, *scrambled* (*Control*) shRNA encoding AAV8-eGFP particles, and *Fpgs* shRNA encoding AAV8-eGFP particles. (C) *Fpgs* transcript level in iWAT, mWAT, gWAT and BAT in control and *Fpgs* kd mice (*n* = 3). (D) Schematic of mouse experiment and timelines. (E-H) Body weight (E), food consumption (F), fat mass (G), and lean mass (H) over time (*n* = 9, Two way ANOVA - multiple comparisons) (J) Liver weight (*n* = 9. Welch’s t test). (J, K) Gene transcript levels for indicated adipogenesis (J) and adipokine (K) marker genes in *Control* and *Fpgs* kd mice iWAT (*n* = 9, Welch’s t test). All bar graphs are represented as mean ± SEM, data points represent biological replicates. Time course plots are represented as mean ± SEM. Exact *P* values are shown in the graphs. Data not marked with exact *P* values are not significant (*P* > 0.05).

**Figure S5.**
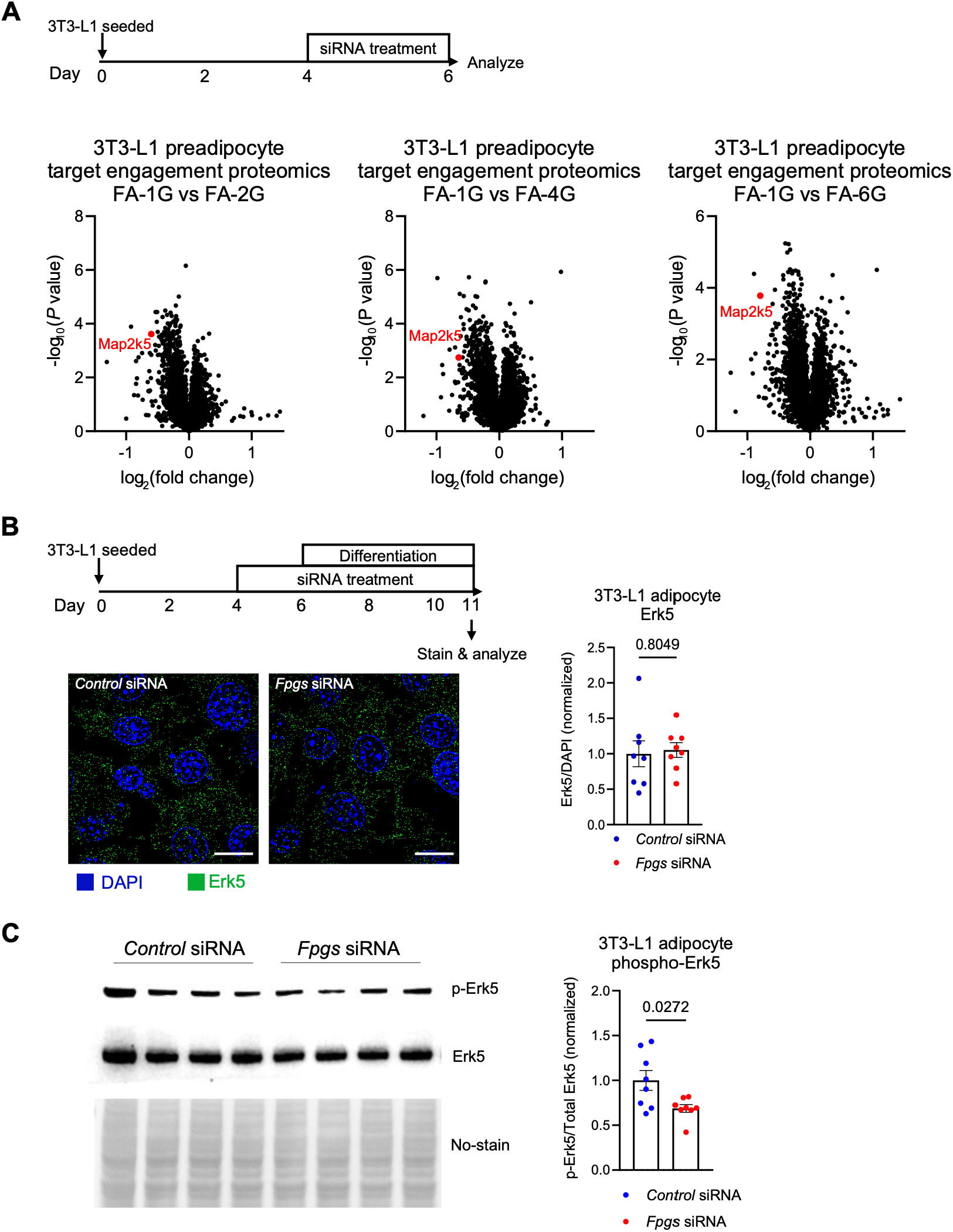
Folic acid monoglutamate binds and inhibits mitogen activated protein kinase kinase 5 (Map2k5). Related to Figure 4. (A) Volcano plots displaying relative thermal shift of proteins when they are bound to monoglutamylated folic acid (FA-1G) compared to when bound to polyglutamylated folic acids. (B) Immunocytochemistry displaying Erk5 (green) and nucleus (blue) in control and *Fpgs* kd 3T3-L1 adipocytes (*n* = 8, Welch’s t test. Scale bar is 10 μm). (C) Western blot for Erk5 and pErk5 in *Control* and *Fpgs* siRNA treated 3T3-L1 adipocytes (*n =* 8, Welch’s t test). All bar graphs are represented as mean ± SEM, data points represent biological replicates.

**Figure S6.**
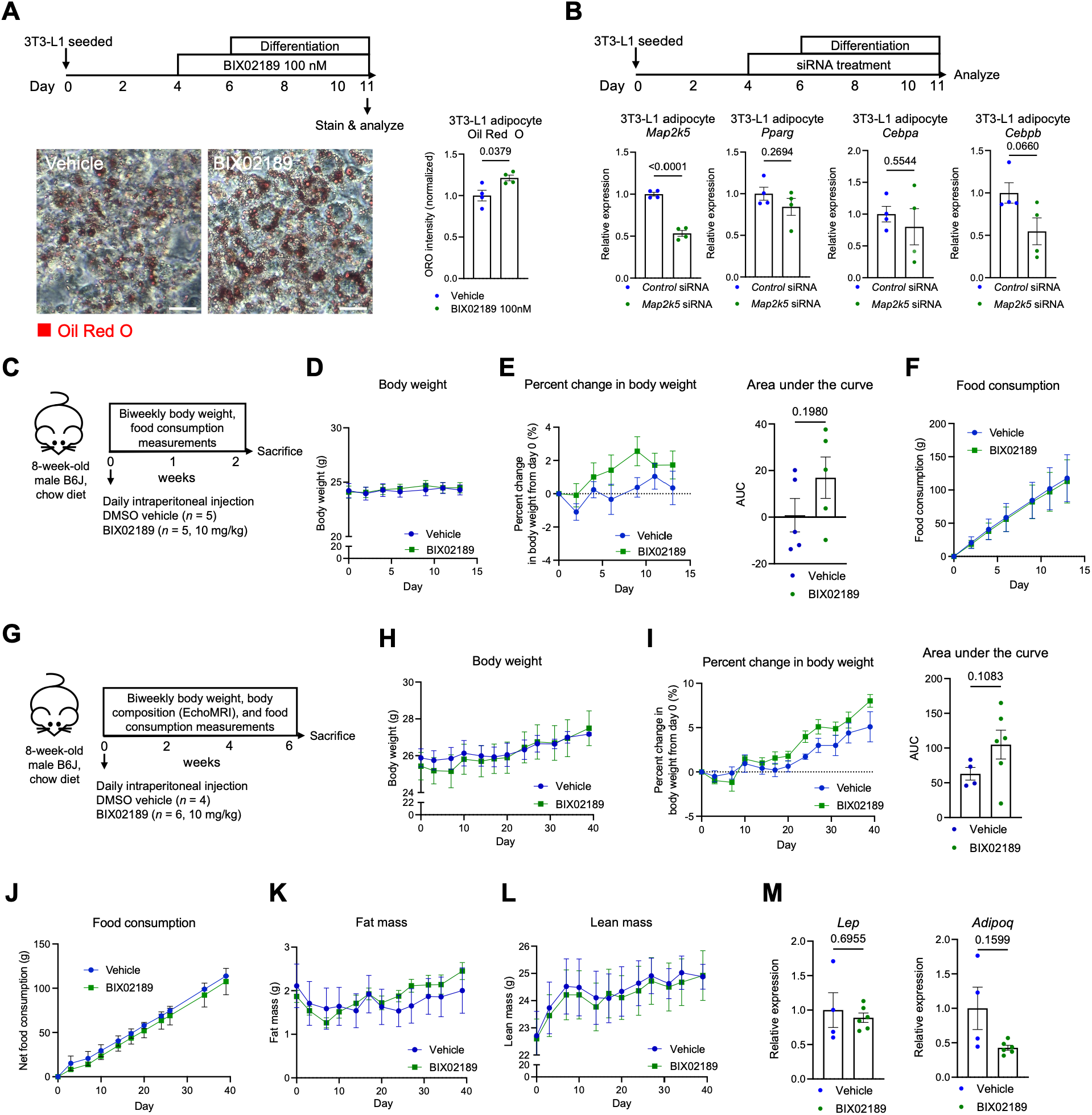
Map2k5 activity promotes lipolytic gene expression to prevent adipocyte hypertrophy. Related to Figure 5. (A) Representative images of vehicle and BIX02189 treated 3T3-L1 adipocytes stained with Oil Red O and quantification (*n* = 4, Welch’s t test. Scale bar is 10 μm). (B) Gene transcript levels in control and *Map2k5* kd 3T3-L1 adipocytes (*n* = 4, Welch’s t test). (C) Schematic of acute BIX02189 treatment mouse experiment and timeline. (D-F) Body weight (D), percent change in body weight with area under the curve (E), food consumption (F), over time (*n* = 5, Two way ANOVA - multiple comparisons, Welch’s ttest for AUC). (G) Schematic of chronic BIX02189 treatment mouse experiment and timeline. (H-L) Body weight (H), percent change in body weight with area under the curve (I) food consumption (J), fat mass (K), and lean mass (L) over time (*n* = 4 for vehicle, *n* = 6 for BIX02189, Two way ANOVA – multiple comparisons, Welch’s t test for AUC). (M) Gene transcript levels in iWAT. All bar graphs are represented as mean ± SEM, data points represent biological replicates. Time course plots are represented as mean ± SEM. Exact *P* values are shown in the graphs. Data not marked with exact *P* values are not significant (*P* > 0.05).

**Table S1.** Analytical parameters of folate extraction method (*n* = 2 technical replicates).

| Vitamer | Limit of Detection (nM) | Linear Range (µM) | Recovery Rate (%) | Run to Run Variation (%) | Retention time (min) | Linear calibration equation | R<sup>2</sup> |
| --- | --- | --- | --- | --- | --- | --- | --- |
| Folic acid | 5 | 10-50 | 83 | 2.68 | 3.64 | Y=1.032E4X | 0.9973 |
| 5-m-THF | 5 | 10-50 | 72 | 8.36 | 3.78 | Y=1.802E3X | 0.9947 |

## STAR★METHODS

### KEY RESOURCES TABLE

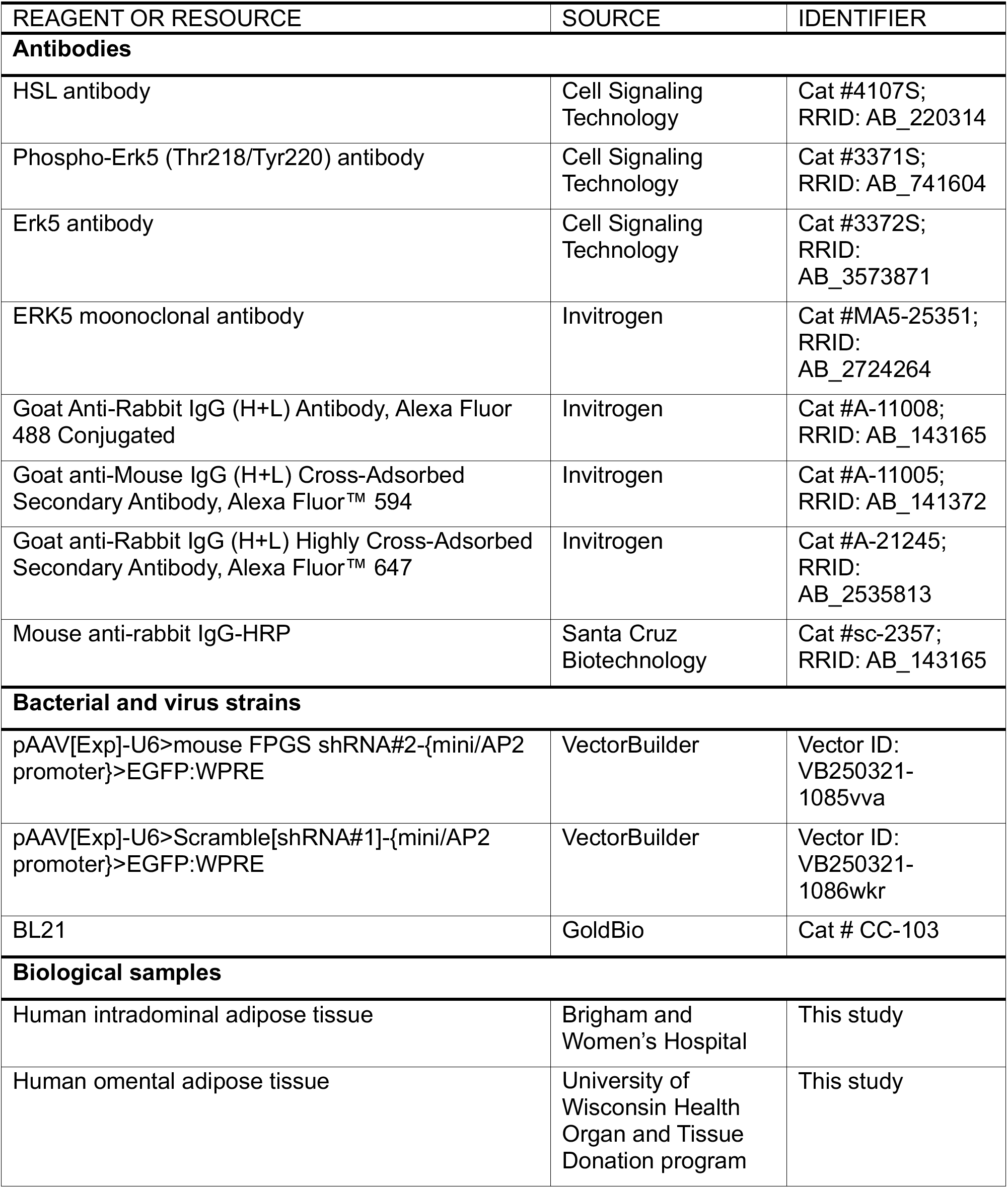
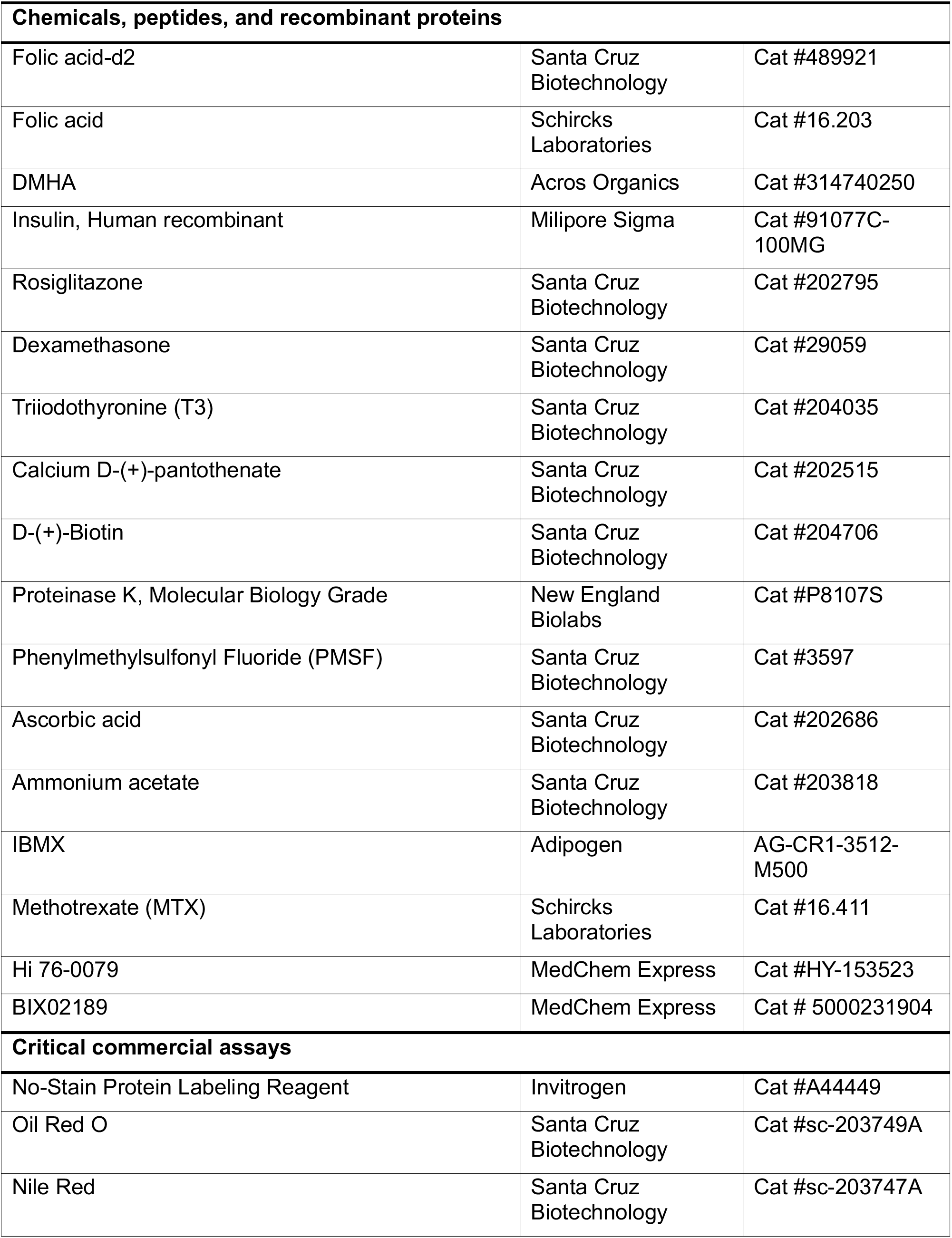
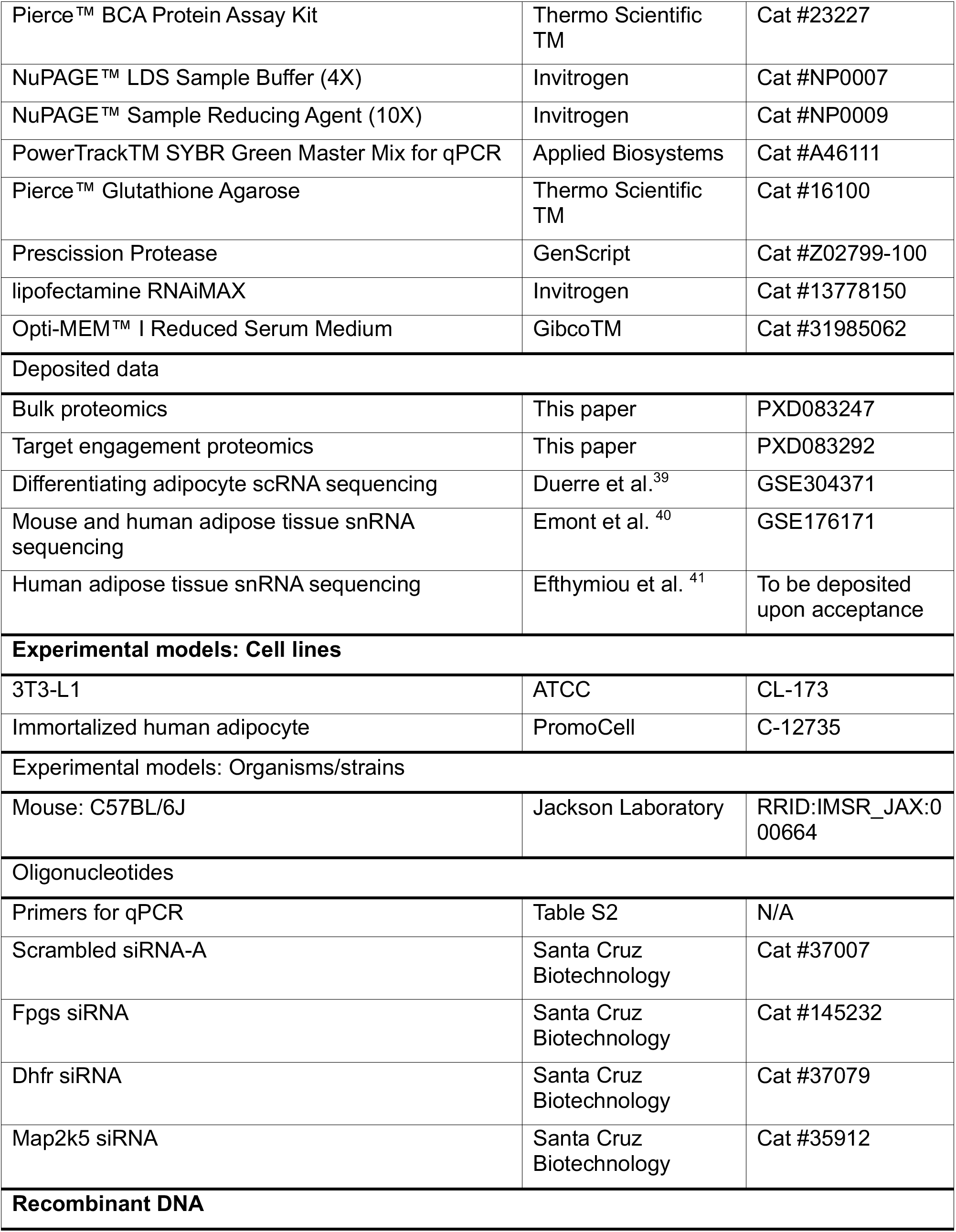
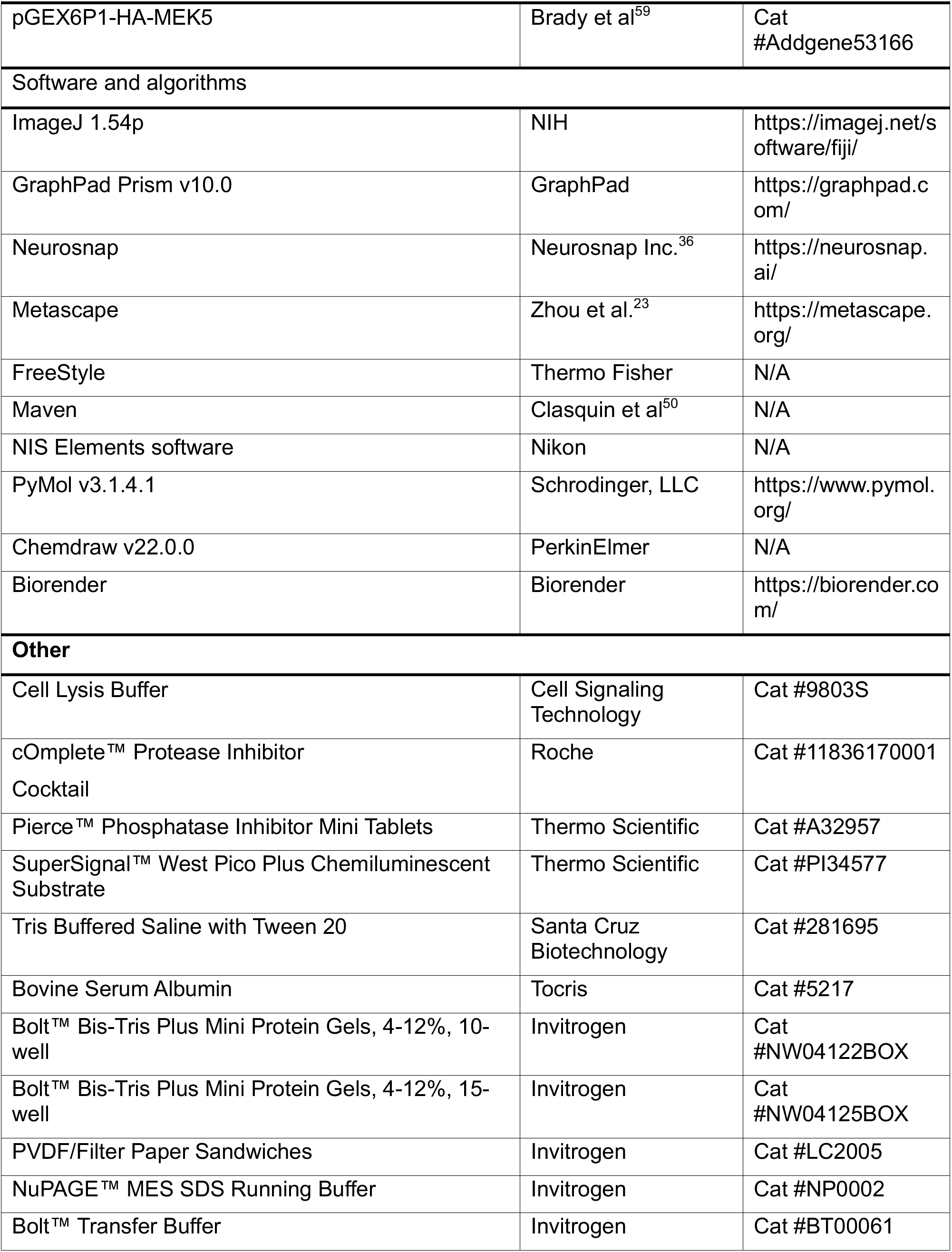
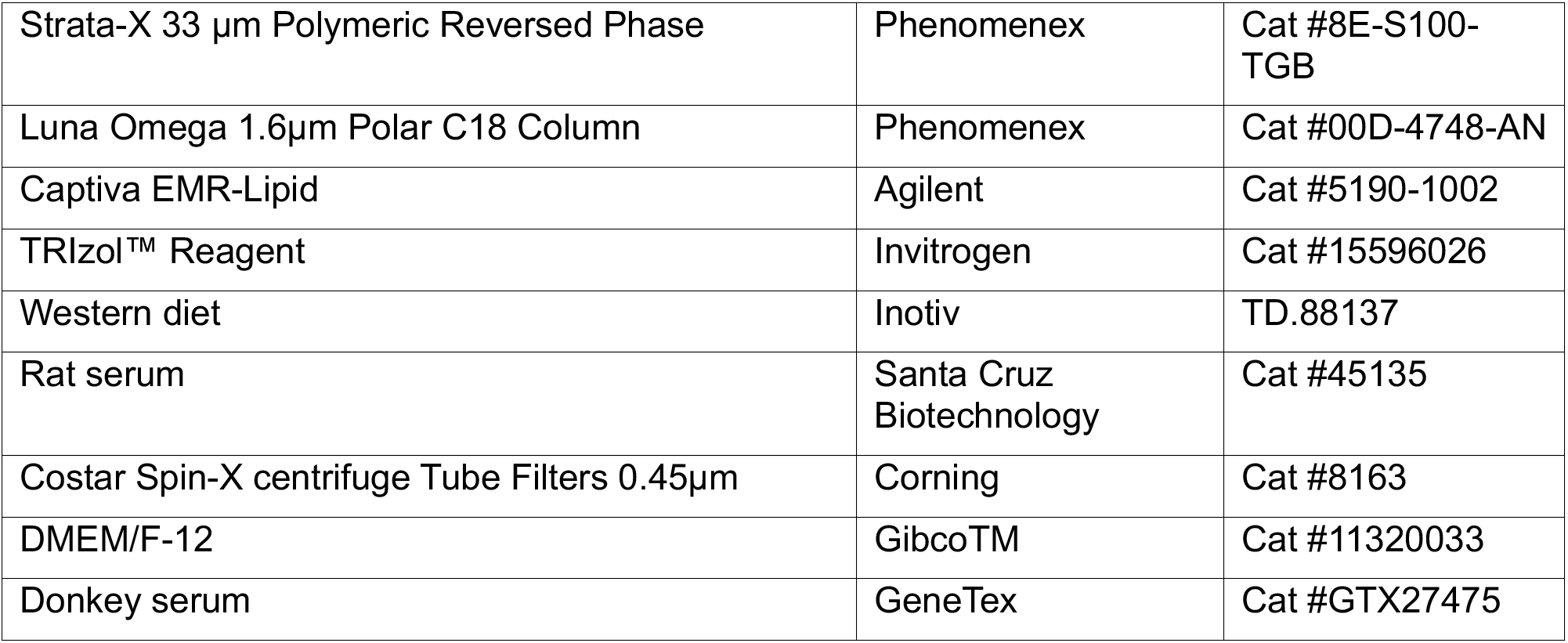

### EXPERIMENTAL MODEL AND STUDY PARTICIPANT DETAILS

#### Mice

For dietary obesity and sleeve gastrectomy (SG) model mice experiments, 5-week-old male, C57BL/6J mice were preconditioned on western diet (WD, 42% calories from fat; 30% w/v sucrose) for 12 weeks to induce obesity. At 17 weeks of age, they were weight matched to receive SG. They were on Recovery Gel Diet (Clear H2O, Westbrook, ME) from -2 to post-operative day 7 and were returned to WD. For SG, mice underwent a 1.5 cm midline laparotomy, ligation of the short gastric vessel, and removal of 75% of the stomach and the entire of the non-glandular section. The gastronomy was closed in two layers with 8-0 permanent monofilament suture. All animals received weight-based saline, norocillin, and buprenorphine (Ethiqua XR; Fidelis Animal Health, North Brunswick, NJ).

For other *in vivo* experiments, C57BL/6J male mice were purchased from Jackson Laboratory (Bar Harbor, ME) and bred in the University of Wisconsin-Madison Biomedical Research Model Services (BRMS) facility. Animals were transferred to the Biochemistry animal facility or Wisconsin Institute for Medical Research animal facility for experiments.

Mice were injected intravenously via the tail vein with 3X10^11^ AAV8 particles (Vector builder) containing either *Control* or *Fpgs* shRNA in 100 µL of saline. For the BIX02189 (MedChem Express) experiments, mice were injected intraperitoneally every day for 2-6 weeks with DMSO or BIX02189 (10 mg/kg) in saline. All animals were housed in climate-controlled animal facilities under a 12 h light/12 h dark cycle with ad libitum access to food and water. Body composition analysis was measured using an EchoMRI (EchoMRI-500-A100-A10TM, EchoMRI LLC). Mice were fasted for 4 h prior to sacrifice. Adipose issues were immediately snap-frozen on dry ice and stored at -80°C until further analysis. Samples for IHC and H&E staining were incubated with 4% paraformaldehyde in PBS for two days, then were paraffin embedded until further processing. All procedures were approved by the Institutional Animal Care and Use Committee (IACUC) at the University of Wisconsin-Madison.

#### Human samples

White adipose tissue biopsies from human subjects undergoing bariatric surgery were collected from intraabdominal adipose tissue during laparoscopic elective surgery at Brigham and Women’s Hospital (BWH, Boston, MA). For the sleeve gastrectomy, intraabdominal adipose tissue biopsies were obtained from the gastrocolic or omental anatomical area. Mesentery adipose samples procured from the UW-Health Human Donor Program were collected during OR. Tissues were snap-frozen in dry ice and kept frozen in −80° C until further processing. The study was approved by the Institutional Review Board at Brigham and Women’s Hospital and at the University of Wisconsin-Madison, and all donors provided written informed consent.

#### Cell culture

3T3-L1 cells from ATCC were a kind gift from the Parks lab at University of Wisconsin-Madison. Cells were cultured in Dulbecco’s Modified Eagle Medium (DMEM, Gibco), supplemented with 10% FBS (GenClone) and 1% penicillin and streptomycin (GenClone) (growth media) in a humidified atmosphere at 37°C with 5% CO_2_. Cells were split every two to three days using 0.25% Trypsin-EDTA (Santa Cruz Biotechnology) and differentiated two to three days after plating. To differentiate 3T3-L1 cells into adipocytes, the cells were cultured in growth media until confluent, then incubated in differentiation medium (500µM IBMX, 1µM dexamethasone, 5µg/mL insulin and 100nM rosiglitazone) for 2 days, and maintained in maintenance medium (5µg/mL insulin and 100nM rosiglitazone) thereafter.

For compound treatments *in vitro*, 3T3-L1 cells were treated with BIX02189 dissolved in ethanol, Hsli, or Methotrexate dissolved in DMSO at specified concentrations. Final ethanol and DMSO concentration were <1%. Vehicle controls received an equivalent concentration of ethanol or DMSO. All treatments were performed in growth medium, differentiation medium or maintenance medium.

Primary human subcutaneous preadipocytes (PromoCell C-12375, Heidelberg, Germany) were seeded at 5x103 cells/cm2 and maintained in Dulbecco’s Modified Eagle’s Medium/F12 (DMEM/F12) supplemented with 10% Fetal Bovine Serum (FBS) and 1% Penicillin/Streptomycin (P/S) with fresh media replacement every 2-3 days. Cells were maintained at 37°C in a humidified 5% CO₂ incubator and monitored daily.

For immortalization, the primary human subcutaneous preadipocytes were transduced with retroviral pBABE-puro-hTERT^50^, which was a gift from Bob Weinberg. To generate retroviral media for transduction, Phoenix-Ampho cells were cultured in DMEM supplemented with 10% FBS and 1% P/S. At 70% confluence, cells were transfected with 10 µg pBABE-puro-hTERT plasmid DNA using JetOPTIUMS transfection buffer and reagent (Polyplus, Illkirch-Graffenstaden, France) per manufacturer’s protocol. Cell culture supernatant was collected after 48 hours of transfection, filtered through a 0.45-µm syringe filter (CellTreat Scientific Products, Pepperell, MA), and stored at -80⁰C. Primary human subcutaneous preadipocytes were transduced with filtered pBABE-puro-hTERT retroviral supernatant and standard growth media in a 1:1 (v/v) ratio with 8 µg/mL Polybrene (Sigma-Aldrich, St. Louis, MO). Cells were transduced in retroviral-conditioned media for 24 hours and then passaged in a 1:2 split ratio. 24 hours later, 1 µg/mL puromycin was added to select for transduced cells.

To differentiate immortalized primary human subcutaneous preadipocytes^51,52^, two days post-confluence, standard growth media was changed to Differentiation Medium 1 for seven days: DMEM/F12 with 3% FBS, 1% P/S, 0.5 mM 3-Isobutyl-1-methylxanthine (IBMX), 0.5 µM dexamethasone, 1 µg/mL insulin, 2 nM Triiodothyronine (T3), 17 µM pantothenate, 33 µM biotin, and 0.5 µM rosiglitazone. On day 7 of differentiation, media was changed to Differentiation Medium 2 for an additional seven days: DMEM/F12 with 3% FBS, 1% P/S, 0.1 µg/mL insulin, and 10 nM dexamethasone. All differentiation media components were purchased from Sigma-Aldrich (St. Louis, MO) except for Rosiglitazone, FBS, and P/S. Media was changed every 2-3 days.

3T3-L1 and human subcutaneous human preadipocytes cells were seeded in 96-, or 12-well plates in growth medium per well and transfected with siRNAs at 100% confluency. Transfection was performed using Lipofectamine RNAiMAX Transfection Reagent. For each well in 96 well plate, 0.6 μL of Lipofectamine RNAiMAX (Invitrogen) and 0.2 μL of 10 μM siRNA (Santa Cruz Biotechnology) were separately diluted in 10 μL Opti-MEM (Gibco), combined at a 1:1 ratio, and incubated for 5 min at room temperature. The resulting 20 μL siRNA–lipid complexes were added to cells and cells were incubated for the indicated duration before downstream analyses. Each well in 12 well plate received 10X the volume of each component.

### METHOD DETAILS

#### Metabolomics

##### Folate extraction

For blood, approximately 50 mg of packed red blood cells were weighed and added to a 2-ml bead-homogenizing tube with glass. Then, 400 μL of folate extraction buffer (20 mM ascorbic acid, 20 mM ammonium acetate and 20 mM β-mercaptoethanol (BME), pH 8.1, adjusted with sodium hydroxide, prepared fresh) was added to each tissue sample and homogenized for 1 min at 5 m s−1 (Bead Ruptor Elite, Omni International). For serum and plasma, 50 μL of sample was pipetted into folate extraction buffer and vortexed for 30 s. For all samples internal standards were spiked into extraction buffer before homogenization. Homogenates were kept on ice unless noted otherwise. Homogenates were centrifuged at 18,200g for 5 min at 4 °C. For monoglutamylated folate analysis, 200 μL of supernatant was aliquoted into a 1.5-ml Eppendorf tube and incubated at 37 °C for 45 min. For total folate analysis, 200 μL of supernatant was aliquoted into a 1.5-ml Eppendorf tube, 5 μL of pooled rat serum was added and sample was vortexed briefly to mix, before incubating at 37 °C for 45 min. After incubation, tubes were stored on ice before being boiled at 100 °C for 5 min. Samples were centrifuged at 18,200g for 5 min to pellet the remaining debris. Supernatants were filtered by SPE using a Strata-X 33-μm polymeric reverse-phase 96-well SPE plate (Phenomenex). After each solvent addition, samples were passed through the column by centrifugation at 3,000g for 30 s. SPE wells were conditioned using 200 μL of methanol and equilibrated using 200 μL of resuspension buffer (20 mM ammonium acetate and 20 mM BME, pH 8.1). Sample supernatant was passed through 0.45 µm filter tubes. Filtered samples were dried down in a SpeedVac (Thermo Scientific, Savant SPD120 and Savant UVS450) at 35 °C for 90 min. Dried down samples were resuspended in 40 μL of resuspension buffer and were moved to autosampler vials (Fisher Scientific, 3452232) for LC-MS analysis. Water for folate extraction was obtained from a GenPure Pro water purification system (Thermo Scientific).

For adipose tissues, approximately 100 mg of tissues were homogenized in 350 µl of folate extraction buffer mixed with 300 µl of ethanol and 50 µl of methanol. For 3T3-L1 preadipocytes, approximately 8 million cells were collected and vortexed with 350 µl of folate extraction buffer mixed with 300 µl of ethanol and 50 µl of methanol. Internal standards were spiked in before homogenization. Homogenates were boiled at 100 °C for 5 min. The samples were then centrifuged at 4 °C at 18,200g for 15 min. The supernatants were filtered through SPE using Captiva EMR (Agilent). The samples were passed through the SPE column using vacuum, followed by elution using 100 µl of 2:1:1 mixture of folate extraction buffer, methanol and ethanol, passed through twice into a 1.5-ml Eppendorf tube. Each sample was divided into half. Next, 5 µl of rat serum was added to one fraction for polyglutamylation analysis. The samples were incubated at 37 °C for 45 min. Samples were next boiled at 100 °C for 5 min, followed by immediately cooling down on ice. Samples were then filtered using Spin-X (Corning) by centrifuging at 18,200g for 5 min. Drying in SpeedVac and subsequent steps were identical to other tissues as described above. For folate extraction from glutathione agarose beads, standard folate extraction as described above was performed, omitting homogenization and rat serum incubation.

#### LC-MS metabolomics

LC-MS metabolomics was performed using a Vanquish ultrahigh-performance LC system (Thermo Scientific) coupled to an Orbitrap Exploris 120 MS instrument (Thermo Scientific). Chromatography was performed using a reverse-phase polar C18 column (Phenomenex). The column chamber and preheater were set to 40 °C. Solvent A was 95% H2O and 5% methanol with 5 mM DMHA (Acros Organics) pH 8.1, adjusted with formic acid. Solvent B was 100% methanol with 5 mM DMHA. All solvents were LC-MS grade purchased from Fisher Chemical (Thermo Scientific). The total run time was 12 min. Flow rate was held constant at 0.4 ml min−1. The chromatography gradient was as follows: 5% solvent B for 0.5 min, linear increase to 85% B over 5.2 min, increase to 100% B for 1.3 min and decrease to 5% to equilibrate the column for 4 min. Eluent from the column was analyzed by MS from the start of the run until 5 min, after which flow was directed to waste for the remainder of the run. Ionization was performed using a HESI source in negative mode at 2,500 V. Nitrogen gas flow was set to 50 (sheath), 10 (auxiliary) and 1 (sweep) in arbitrary units. The ion transfer tube was maintained at 325 °C with the vaporizer at 350 °C. Quantification was performed using Freestyle version 1.8.65.0 (Thermo Fisher) and MAVEN78 version 2.10.21 to identify and calculate integrated peak areas. Manual curation confirmed peak identification and parameters. All signals were normalized to internal standards (methotrexate, folic acid-d2) for injection variation and compared to the standard curve before reporting molar concentrations normalized to sample mass. The standard curve was modeled using linear regression. Total measured folate in each sample was calculated by adding all detected and measured folate vitamers. Standard curves were prepared with folate standards purchased from Schircks Laboratories. Standard stocks were made at 200 µM and serially diluted until signal could not be detected to determine limit of detection and linear range. Internal standards purchased from Schircks Laboratories (methotrexate,) and Santa Cruz Biotechnology (folic acid-d2,).

#### Lipid staining

3T3-L1 preadipocytes were stained with 3µM Nile Red (SCBT) in the growth media for 30 minutes at 37 °C, washed with PBS, and fixed with 4% paraformaldehyde in PBS for 10 minutes at room temperature. Nuclei was counter stained with DAPI (1:3000 in PBS) for 10 minutes at room temperature. Fluorescence images were obtained using Nikon A1R-Si+ Confocal Microscope. 3T3-L1 adipocytes were washed with PBS, fixed with 4% paraformaldehyde in PBS for 10 minutes at room temperature, washed with 60% isopropanol, incubated with 0.2% Oil Red O (SCBT) staining in 60% isopropanol for 5 minutes, and washed with deionized water until being imaged. Total lipid contents in 3T3-L1 adipocytes were quantified by dissolving Oil Red O stains in 60% isopropanol and measuring absorbance at 518 nm. Images were obtained using Zeiss primovert microscope.

#### Electron Microscopy

3T3-L1 preadipocytes were processed and imaged UW-Madison Electron Microscopy Facility. Briefly, cells were fixed in a modified Karnovsky’s fixative (2.5% glutaraldehyde/2.0% formaldehyde in 0.1 M sodium phosphate buffer (PB), pH 7.2), post fixed in 1.0% osmium tetroxide in 0.1M PB, dehydrated in a grade series of ethanol, transitioned in acetone, and embedded in EmBed 812 (Electron Microscopy Sciences). Polymerized samples were sectioned (80nm) on an ultramicrotome and sections were placed onto formvar coated 2x1 Cu slot girds and post-stained with uranyl acetate and lead citrate. The sections were viewed at 80kV on a FEI CM120 TEM (Thermo Fisher Scientific) and imaged on an AMT BioSprint 12 digital camera (AMT Imaging)

#### Untargeted bulk proteomics

##### Sample preparation for mass spectrometry

Samples for protein analysis were prepared essentially as previously^53^. 3T3-L1 cells were washed three times with ice-cold PBS. 100 µL of lysis buffer (8 M urea, 200 mM EPPS pH 8.5, cOmplete™ Protease Inhibitor Cocktail (Roche), 0.1% SDS) was added and shaken for 10 min at 4°C to allow cell lysis. 100 µL of nuclease buffer (200 mM EPPS pH 8.5, 0.5 µL/mL Pierce™ Universal Nuclease (Thermo Fisher)) was then added.

Following lysis, 25 μg of protein from each sample was reduced with tris(2-carboxyethyl)phosphine (TCEP) (Thermo Scientific, #77720), alkylated with iodoacetamide (MP Biochemicals, #100351), and then further reduced with dithiothreitol (DTT) (Alfa Aesar, #A15797). A buffer exchange was carried out using a modified SP3 protocol. Briefly, ∼250 µg of Cytiva SpeedBead Magnetic Carboxylate Modified Particles (65152105050250 and 4515210505250), mixed at a 1:1 ratio, were added to each sample. 100% ethanol was added to each sample to achieve a final ethanol concentration of at least 50%. Samples were incubated with gentle shaking for 15 min. Samples were washed three times with 80% ethanol. Protein was eluted from SP3 beads using 200 mM EPPS pH 8.5 containing Lys-C (Wako, #129-02541). Samples were digested overnight at room temperature with vigorous shaking. The next morning trypsin (1:50 enzyme to protein ratio) (Thermo Scientific, #90305) was added to each sample and further incubated for 6 h at 37° C. Acetonitrile was added to each sample to achieve a final concentration of ∼33%. Each sample was labelled, in the presence of SP3 beads, with ∼62.5 µg of TMTPro reagents (Thermo Fisher Scientific). Following confirmation of satisfactory labelling (>97%), excess TMT was quenched by addition of hydroxylamine to a final concentration of 0.3%. The full volume from each sample was pooled and acetonitrile was removed by vacuum centrifugation for 1 h. The pooled sample was acidified, and peptides were de-salted using a Sep-Pak 50mg tC18 cartridge (Waters). Peptides were eluted in 70% acetonitrile, 1% formic acid and dried by vacuum centrifugation.

##### Basic pH reversed-phase separation (BPRP)

TMT labeled peptides were solubilized in 5% acetonitrile/10 mM ammonium bicarbonate, pH 8.0 and ∼300 µg of TMT labeled peptides were separated by an Agilent 300 Extend C18 column (3.5 µm particles, 4.6 mm ID and 250 mm in length). An Agilent 1260 binary pump coupled with a photodiode array (PDA) detector (Thermo Scientific) was used to separate the peptides. A 45-minute linear gradient from 10% to 40% acetonitrile in 10 mM ammonium bicarbonate pH 8.0 (flow rate of 0.6 mL/min) separated the peptide mixtures into a total of 96 fractions (36 seconds). A total of 96 fractions were consolidated into 24 samples in a checkerboard fashion and vacuum dried to completion. Each sample was desalted via Stage Tips and re-dissolved in 5% formic acid/ 5% acetonitrile for LC-MS3 analysis.

##### Liquid chromatography separation and tandem mass spectrometry (LC-MS3)

Proteome data were collected on an Orbitrap Eclipse mass spectrometer (Thermo Fisher Scientific) coupled to a Proxeon EASY-nLC 1000 LC pump (Thermo Fisher Scientific). A FAIMS device was enabled during data collection with CV values set to -40, -60, and -80. Fractionated peptides were separated using a 120 min gradient at 500 nL/min on a 35 cm column (i.d. 100 μm, Accucore, 2.6 μm, 150 Å) packed in-house. MS1 data were collected in the Orbitrap (60,000 resolution; maximum injection time 50 ms; AGC 4 × 105). Charge states between 2 and 5 were required for MS2 analysis in the ion trap, and a 120 second dynamic exclusion window was used. Top 10 MS2 scans were performed in the ion trap with CID fragmentation (isolation window 0.5 Da; Turbo; NCE 35%; maximum injection time 35 ms; AGC 1 × 104). Real-time search was used to trigger MS3 scans for quantification. MS3 scans were collected in the Orbitrap using a resolution of 50,000, NCE of 55%, maximum injection time of 250 ms, and AGC of 1.25 × 105. The close out was set at two peptides per protein per fraction.

##### Data analysis

Raw files were converted to mzXML, and monoisotopic peaks were re-assigned using Monocle^54^. Searches were performed using the Comet search algorithm against a mouse database (May 2025) downloaded from Uniprot. A 50 ppm precursor ion tolerance, 1.0005 fragment ion tolerance, and 0.4 fragment bin offset was used for MS2 scans collected in the ion trap. TMTpro on lysine residues and peptide N-termini (+304.2071 Da) and carbamidomethylation of cysteine residues (+57.0215 Da) were set as static modifications, while oxidation of methionine residues (+15.9949 Da) was set as a variable modification.

Each run was filtered separately to 1% False Discovery Rate (FDR) on the peptide-spectrum match (PSM) level. Then proteins were filtered to the target 1% FDR level across the entire combined data set. For reporter ion quantification, a 0.003 Da window around the theoretical m/z of each reporter ion was scanned, and the most intense m/z was used. Reporter ion intensities were adjusted to correct for isotopic impurities of the different TMTpro reagents according to manufacturer specifications. Peptides were filtered to include only those with a summed signal-to-noise (SN) ≥ 180 across all TMT channels. For each protein, the filtered peptide TMTpro SN values were summed to generate protein or phosphorylation site quantification values. The signal-to-noise (S/N) measurements of peptides assigned to each protein were summed (for a given protein). These values were normalized so that the sum of the signal for all proteins in each channel was equivalent thereby accounting for equal protein loading, and *p* values were adjusted for multiple comparisons using the Benjamini-Hochberg (BH) correction.

##### Functional enrichment analysis

Gene Ontology (GO) and pathway enrichment analyses were performed using Metascape^23^. Differentially expressed proteins (cutoff: fold-change > 1.75 or < 1.75 and Benjamini-Hochberg adjusted *p* < 0.05) were used for enrichment analysis. GO pathway analyses were performed using default parameters.

#### Target Engagement Proteomics

##### Lysate-based proteome integral solubility alteration

Frozen 3T3-L1 cell pellets were thawed on ice and resuspended in lysis buffer (1 X PBS pH 7.4, 1 mM MgCl_2_, protease inhibitor). The proteomes were extracted using a dounce homogenizer (20 strokes). The extracts were spun at 300 x *g* for 3 min to remove any unbroken cells. The resulting crude extract was diluted to 2 mg/mL in lysis buffer. Each compound was added to lysis buffer at a 2 X concentration. In order to initiate the experiment, an equal volume of crude extract and treatment buffer were combined, to achieve a final protein concentration of 1 mg/mL and compound concentration of 10 μM, and incubated for 30 min. After incubation, an equal volume of each sample was transferred to 10 PCR tubes. The PCR tubes were heated across a thermal gradient ranging from 48°C to 58°C for 3 min to induce thermal denaturation. An equal volume from each PCR tube was pooled. An equal volume of extraction buffer (1 X PBS pH 7.4, 1% NP-40, protease inhibitors) was added to added to each pooled sample to achieve a final NP-40 concentration of 0.5%. Samples were incubated for 10 min at 4°C on a roller. Extracted samples were spun at 21,000 x *g* for 90 min to separate insoluble aggregates from soluble protein. An equal volume from each soluble fraction was collected and prepared for LC-MS/MS analysis.

##### LC-MS/MS sample preparation

Samples (15-20 μg protein) were diluted in prep buffer (400 mM EPPS pH 8.5, 1% SDS, 10 mM tris(2-carboxyethyl)phosphine hydrochloride) and incubated at room temperature for 10 min. Iodoacetimide was added to a final concentration of 10 mM to each sample and incubated for 25 min. in the dark. Finally, DTT was added to each sample to a final concentration of 10 mM. A buffer exchange was carried out using a modified SP3 protocol. Briefly, ∼250 μg of each SpeedBead Magnetic Carboxylate modified particles (Cytiva; 45152105050250, 65152105050250) mixed at a 1:1 ratio were added to each sample. 100% ethanol was added to each sample to achieve a final ethanol concentration of at least 50%. Samples were incubated with gentle shaking for 15 min. Samples were washed three times with 80% ethanol. Protein was eluted from SP3 beads using 200 mM EPPS pH 8.5 containing trypsin (ThermoFisher Scientific) and Lys-C (Wako). Samples were digested overnight at 37°C with vigorous shaking. Acetonitrile was added to each sample to achieve a final concentration of 30%. Each sample was labelled, in the presence of SP3 beads, with ∼65 μg of TMTpro-16plex reagents^55^ (ThermoFisher Scientific). Experimental layouts for each experiment were described in corresponding source data tables. Following confirmation of satisfactory labelling (>97%), excess TMTpro reagents were quenched by addition of hydroxylamine to a final concentration of 0.3%. The full volume from each sample was pooled and acetonitrile was removed by vacuum centrifugation for one hour. The pooled sample was acidified using formic acid and peptides were de-salted using a Sep-Pak Vac 50 mg tC18 cartridge (Waters). Peptides were eluted in 70% acetonitrile, 1% formic acid and dried by vacuum centrifugation. The peptides were resuspended in 10 mM ammonium bicarbonate pH 8, 5% acetonitrile and fractionated by basic pH reverse phase HPLC. In total 24 fractions were collected. The fractions were dried in a vacuum centrifuge, resuspended in 5% acetonitrile, 1% formic acid and desalted by stage-tip. Final peptides were eluted in, 70% acetonitrile, 1% formic acid, dried, and finally resuspended in 5% acetonitrile, 5% formic acid. In the end, 12 of 24 fractions were analyzed by LC-MS/MS.

##### Mass spectrometry data acquisition

Data were collected on an Orbitrap Eclipse mass spectrometer (ThermoFisher Scientific) coupled to a Proxeon EASY-nLC 1000 LC pump (ThermoFisher Scientific). Peptides were separated using a 120-min gradient at 500 nL/min on a 30-cm column (i.d. 100 μm, Accucore, 2.6 μm, 150 Å) packed in house. High-field asymmetric-waveform ion mobility spectroscopy (FAIMS) was enabled during data acquisition with compensation voltages (CVs) set as −40 V, −60 V, and −80 V. MS1 data were collected using the Orbitrap (60,000 resolution; maximum injection time 50 ms; AGC 4 × 10^5^). Determined charge states between 2 and 6 were required for sequencing, and a 60 s dynamic exclusion window was used. Data dependent mode was set as cycle time (1 s). MS2 scans were performed in the Orbitrap with HCD fragmentation (isolation window 0.5 Da; 50,000 resolution; NCE 36%; maximum injection time 86 ms; AGC 1 × 10^5^).

##### Mass spectrometry data analysis

Raw files were first converted to mzXML, and monoisotopic peaks were re-assigned using Monocle. Database searching included all human entries from Uniprot (downloaded in February 2020). The database was concatenated with one composed of all protein sequences in the reversed order. Sequences of common contaminant proteins (e.g., trypsin, keratins, etc.) were appended as well. Searches were performed using the comet search algorithm. Searches were performed using a 50-ppm precursor ion tolerance and 0.02 Da product ion tolerance. TMTpro on lysine residues and peptide N termini (+304.2071 Da) and carbamidomethylation of cysteine residues (+57.0215 Da) were set as static modifications, while oxidation of methionine residues (+15.9949 Da) was set as a variable modification.

Peptide-spectrum matches (PSMs) were adjusted to a 1% false discovery rate (FDR). PSM filtering was performed using linear discriminant analysis (LDA) as described previously^56^ while considering the following parameters: comet log expect, different sequence delta comet log expect (percent difference between the first hit and the next hit with a different peptide sequence), missed cleavages, peptide length, charge state, precursor mass accuracy, and fraction of ions matched. Each run was filtered separately. Protein-level FDR was subsequently estimated at a data set level. For each protein across all samples, the posterior probabilities reported by the LDA model for each peptide were multiplied to give a protein-level probability estimate. Using the Picked FDR method^57^, proteins were filtered to the target 1% FDR level.

For reporter ion quantification, a 0.003 Da window around the theoretical *m/z* of each reporter ion was scanned, and the most intense *m/z* was used. Reporter ion intensities were adjusted to correct for the isotopic impurities of the different TMTpro reagents according to manufacturer specifications. Peptides were filtered to include only those with a summed signal-to-noise (SN) of 160 or greater across all channels. For each protein, the filtered peptide TMTpro SN values were summed to generate protein quantification.

#### Molecular analyses

##### RT-PCR

RNA extraction and cDNA synthesis was done as previously described^18^. qPCR was performed using PowerTrack SYBR Green Master Mix (Invitrogen) in a 384-well format. C*_q_* values were determined using QuantStudio 7 Pro (Thermo Fisher). The 2^−ΔΔ*Ct*^ method was used to calculate the relative change in gene expression with the murine *ppib* gene used as the loading control. Primers were designed using NIH Primer-BLAST and purchased from Integrated DNA Technologies (Coralville, IA).

##### Western Blotting

3T3-L1 adipocytes were lysed in 1x Cell Lysis Buffer (Cell Signaling Technology) supplemented with a protease (Roche) and phosphatase inhibitor cocktail (Thermo Fisher). Samples were centrifuged at 18,00 x *g* for 10 min at 4°C, and supernatants were used for downstream applications. Protein concentrations were determined using a BCA assay (Thermo fisher) with bovine serum albumin (BSA) as a standard. Proteins were mixed with 4x loading buffer (Invitrogen) and 10x reducing agent (Invitrogen), boiled at 95°C for 5 min, and run on a precast Bolt Bis-Tris SDS-PAGE gel (Invitrogen). For p-Erk5, samples were incubated at room temperature for 5 min before being run. Proteins were transferred onto PVDF membranes using wet transfer at 30 V for 1 h. Total protein was visualized using No-Stain Protein Labeling Reagent (Invitrogen) according to the manufacturer’s protocol. Membranes were then blocked in 5% nonfat milk (Cell Signaling Technology) in TBST for 1 h at room temperature. Primary antibody incubations were done overnight at 4°C for Hsl and total Erk5 and for two days for p-Erk5 with antibodies diluted 5% nonfat milk in TBST. After washing with TBST, membranes were incubated with HRP-conjugated secondary antibodies diluted in 5% nonfat milk in TBST for 1 h at room temperature. Blots were developed using SuperSignal™ West Pico PLUS Chemiluminescent Substrate (Thermo Scientific) and imaged on the iBright imaging system (Invitrogen). Band intensities were quantified using the iBright imaging software. Primary antibodies used include: HSL (4107s, 1:1000), p-ERK5 (3371s, 1:250), ERK5 (3372s, 1:1000). Mouse anti-rabbit IgG-HRP (Santa Cruz, 2357, 1:3000) was used as a secondary antibody.

#### H&E histology analyses

Formalin-fixed and paraffin-embedded adipose tissues were sectioned at 5 µm thickness and dried at 60°C for 20 min prior to staining with hematoxylin and eosin (H&E), p-Erk5, or Erk5 using the Leica ST5010 Autostainer XL (Leica Biosystems) according to the UWCC Experimental Animal Pathology Laboratory protocol. H&E sections were sequentially treated with xylene, graded ethanol, Hematoxylin MX 560, Define MX, Blue Buffer 8, and Eosin 515 Trichrome, followed by dehydration through ethanol and clearing in xylene. p-Erk5 and Erk5 slides were HIER (heat induced antigen retrieval) in pH 6.0 citrate buffer (10 mM citric acid, 0.05% tween 20) for 3 minutes in Decloaking Chamber (Biocare Medical, Concord, CA). Slides were then cooled for 30 min, rinsed in 1x PBS, blocked with 10% goat serum (Sigma, St. Louis, MO) in PBS for 1 hour at room temperature, followed by primary antibody (p-Erk5, 1:200) (Erk5, 1:400) incubation in PBS with 1% goat serum and 0.1% Triton X-100 overnight at 4°C. The slides were then rinsed x 3 with 1x PBS and incubated with Alexafluor 647 goat anti-rabbit and Alexafluor 594 goat anti-mouse at 1:1000 in PBS for 1 hr at room temperature. After the secondary incubation, slides were rinsed x 3 with 1x PBS followed by a DH_2_O rinse, and mounted with a cover slip using Prolong Gold with DAPI (Thermo Fisher Scientific, P36931). Images were acquired with a Nikon A1R+ confocal microscope.

#### Binding Affinity assay

##### Protein purification

GST-tagged human MAP2K5 plasmid (Addgene) was transfected into chemically competent BL21 cell line, which was a generous gift from the Wei lab at University of Wisconsin-Madison. After overnight incubation with 0.1 mM IPTG, cells were pelleted at 3000 g for 15 min. Cell pellet was lysed with bacterial lysis buffer (50 mM Tris-HCl, 150 mM NaCl, 1 mM DTT, 10% glycerol, protease inhibitor cocktail tablet, pH = 8). Supernatant was collected after centrifuging at 3000 g for 15 min. at 4°C. Protein purification was performed according to the manufacturer’s instructions (Thermo fisher). Briefly, after incubating with Glutathione agarose beads at 4°C for 2 hours, the beads were washed and cleaved with GST protease (GenScript).

##### Limited proteolysis assay

To assess the binding affinity of FA-1G to MAP2K5, a limited proteolysis assay was conducted. Reaction buffer consisted of 150 mM NaCl, 50 mM Tris-HCl, 1 mM CaCl2 in pH = 8. All protein and FA stocks were diluted in reaction buffer, and all reactions were set up on ice.

Briefly, 3.5 µg of purified MAP2K5 protein was incubated at 37°C for 30 min. with or without FA (200 µM). Then, 20 ng of Proteinase K was added, and the reaction proceeded for the following time points: t = 0, 2.5 and 5 minutes. Each time point was a separate tube to ensure no change in total protein or folate species. To quench each reaction, 5 mM of phenylmethylsulfonyl fluoride (PMSF, Santa Cruz Biotechnology) was added to the reaction and incubated at room temperature for 10 min. Then, 4x loading buffer and 10x reducing agent (Invitrogen) was added, and the reaction was boiled at 95°C for 5 min and placed on ice until all reactions were complete and quenched with PMSF. After the completion of all reactions, the samples were run on a precast Bolt Bis-Tris SDS-PAGE gel (Invitrogen). Proteins were transferred onto PVDF membranes using wet transfer at 30 V for 1 h, and total protein was visualized on the membrane using No-Stain Protein Labeling Reagent (Invitrogen) according to the manufacturer’s protocol.

##### On-resin ligand binding assay

GST tagged MAP2K5 was allowed to bind to glutathione agarose beads at 4°C for one hour in equilibrium buffer (150 mM NaCl, 50 mM Tris-HCl pH = 8). The beads were then washed three times with ice cold equilibrium buffer, then washed with ligand buffer (10 mM ammonium acetate, 20M NaCl, pH = 8) three times. Beads with protein were then incubated with 200 µM folic acid (Schircks Laboratories) for 30 minutes at 37°C in ligand buffer. The beads were washed three times with ice cold ligand buffer. Folate extraction and LC-MS analysis was performed on the beads as described above.

#### In silico docking

Molecular docking simulations were performed using DiffDock-L on the Neurosnap web platform^36^. The computed structure of human Mitogen Activated Protein Kinase Kinase 5 (MAP2K5, AlphaFold DB: Q13163) was obtained from the Protein Data Bank and used as the receptor structure. Ligand structures in SMILES format were submitted to DiffDock-L using default parameters. Structural visualization and figure rendering were performed using PyMOL v3.1.4.1.

#### Immunocytochemistry

Cells grown on 96-well glass-bottom plates were fixed in 4% formaldehyde and washed three times with PBS (5 minutes per wash). Cells were permeabilized and blocked in PBS containing 4% donkey serum (GeneTex) and 0.3% Triton X-100 for 1 hour at room temperature with gentle agitation. Cells were incubated with primary antibodies diluted in bocking solution overnight at 4°C with gentle agitation. The following day, cells were washed three times in PBS (5 minutes per wash) at room temperature and incubated with fluorophore conjugated secondary antibodies diluted in blocking solution for 1 hour at room temperature, protected from light. After washing with PBS, nuclei were counter stained with DAPI (1:3000 in PBS) for 5 minutes. Cells were washed again with PBS before imaging. Images were obtained using appropriate filter sets on a Nikon A1R+ confocal microscope (Nikon instruments) at the Biochemistry Optical Core (BOC) Facility at UW-Madison. Primary antibodies used for immunocytochemistry fluorescence staining are: phospho-Erk5 (1:250), Erk5 (1:500). Goat anti-rabbit IgG (H+L) Cross-adsorbed Alexa Fluor^TM^ 488 (1:500) was used as a secondary antibody.

**Table S2.**
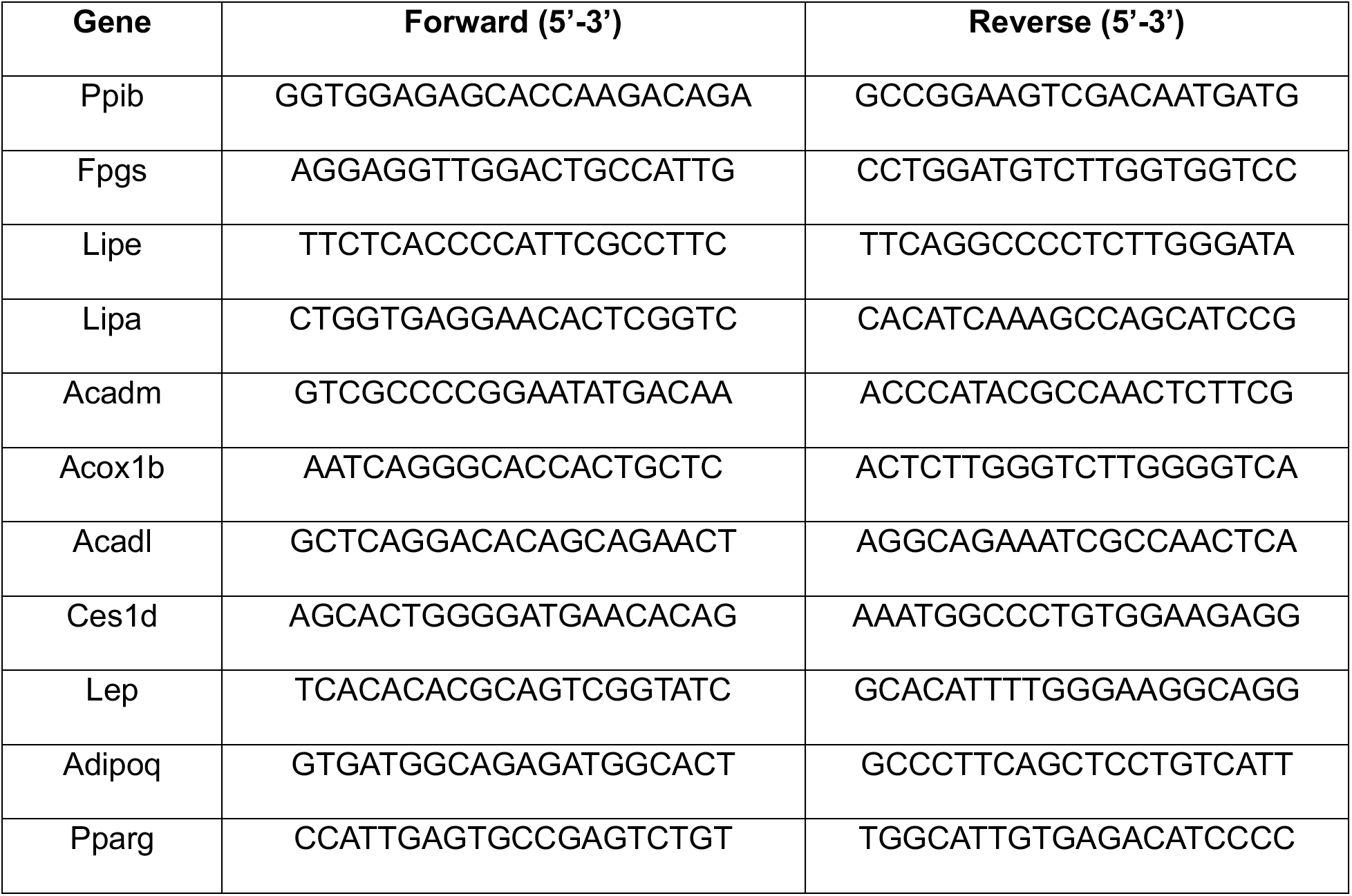
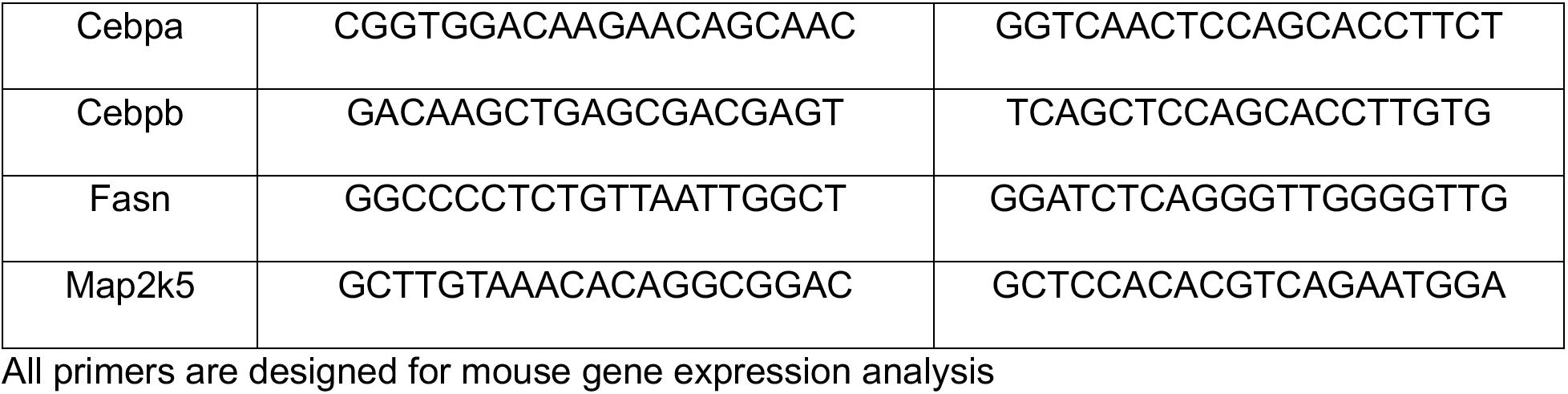
qPCR primer sequences.

#### Graphics

The schematic illustrations were created using BioRender. Chemical structures were generated using ChemDraw. Molecular docking structures and structural visualizations were rendered using PyMOL.

### QUANTIFICATION AND STATISTICAL ANALYSIS

Data are presented as mean ± SEM for bar graphs and scatter dot plots, unless otherwise specified. Violin plots depict data distribution, with the central line indicating the median and lines representing all data points. Each data point represents an individual biological replicate, unless otherwise specified, as defined in the figure legends. For comparisons between two groups, an unpaired two-tailed Welch’s t test was used. For non-normally distributed data, the Mann–Whitney test was applied. For comparisons among three or more groups, one-way ANOVA followed by appropriate post hoc multiple comparison tests was used. When normality assumptions were not met, the Kruskal–Wallis test was used. For experiments involving two independent variables, two-way ANOVA was used for normally distributed data. When multiple parameters were compared between two groups and normality were not met, multiple Mann-Whitney tests were performed. For all two-way ANOVA analyses, a Sidak test was used. For proteomics datasets, *p* values were adjusted for multiple comparisons using the Benjamini-Hoechberg (BH) correction. For publicly available datasets, adjusted *p* values were used for analyses. A power calculation was used to determine total number of animals required for each study. Briefly, a dichotomous endpoint for two independent study groups with an expected error in the mean of 10% and a difference of at least 75% was considered. A false positive rate of 10% and a false negative rate of 20% (power) identified a sample size of at least n = 4 for each study group. A *p* value < 0.05 was considered statistically significant. Exact *p* values are reported in the figures. Data not marked are not statistically significant (*p* > 0.05) or as indicated in each figure legend. Statistical analysis was performed using GraphPad PRISM. Exact *n* values and statistical tests used are listed in the figure legends.

## REFERENCES

1 Stern, J. H. & Scherer, P. E. Adipose tissue biology in 2014: Advances in our understanding of adipose tissue homeostasis. Nat Rev Endocrinol 11, 71–72 (2015). 10.1038/nrendo.2014.219

2 Morigny, P., Boucher, J., Arner, P. & Langin, D. Lipid and glucose metabolism in white adipocytes: pathways, dysfunction and therapeutics. Nat Rev Endocrinol 17, 276–295 (2021). 10.1038/s41574-021-00471-8

3 Jeon, Y. G., Kim, Y. Y., Lee, G. & Kim, J. B. Physiological and pathological roles of lipogenesis. Nat Metab 5, 735–759 (2023). 10.1038/s42255-023-00786-y

4 Heymsfield, S. B. & Wadden, T. A. Mechanisms, Pathophysiology, and Management of Obesity. N Engl J Med 376, 254–266 (2017). 10.1056/NEJMra1514009

5 Sakers, A., De Siqueira, M. K., Seale, P. & Villanueva, C. J. Adipose-tissue plasticity in health and disease. Cell 185, 419–446 (2022). 10.1016/j.cell.2021.12.016

6 Galasso, M. et al. The Impact of Different Nutritional Approaches on Body Composition in People Living with Obesity. Curr Obes Rep 14, 45 (2025). 10.1007/s13679-025-00636-w

7 Hariri, N. & Thibault, L. High-fat diet-induced obesity in animal models. Nutr Res Rev 23, 270–299 (2010). 10.1017/S0954422410000168

8 Suárez, R. et al. Obesity and nutritional strategies: advancing prevention and management through evidence-based approaches. Food and Agricultural Immunology 36 (2025). 10.1080/09540105.2025.2491597

9 Ducker, G. S. & Rabinowitz, J. D. One-Carbon Metabolism in Health and Disease. Cell Metab 25, 27–42 (2017). 10.1016/j.cmet.2016.08.009

10 Chan, C. W., Chan, P. H. & Lin, B. F. Folate Deficiency Increased Lipid Accumulation and Leptin Production of Adipocytes. Front Nutr 9, 852451 (2022). 10.3389/fnut.2022.852451

11 Chen, S. et al. Suppression of high-fat-diet-induced obesity in mice by dietary folic acid supplementation is linked to changes in gut microbiota. Eur J Nutr 61, 2015–2031 (2022). 10.1007/s00394-021-02769-9

12 Zhao, M. et al. Chronic folate deficiency induces glucose and lipid metabolism disorders and subsequent cognitive dysfunction in mice. PLoS One 13, e0202910 (2018). 10.1371/journal.pone.0202910

13 Kelly, K. B. et al. Excess Folic Acid Increases Lipid Storage, Weight Gain, and Adipose Tissue Inflammation in High Fat Diet-Fed Rats. Nutrients 8 (2016). 10.3390/nu8100594

14 Bird, J. K. et al. Obesity is associated with increased red blood cell folate despite lower dietary intakes and serum concentrations. J Nutr 145, 79–86 (2015). 10.3945/jn.114.199117

15 Li, Z. et al. Folate and vitamin B12 status is associated with insulin resistance and metabolic syndrome in morbid obesity. Clin Nutr 37, 1700–1706 (2018). 10.1016/j.clnu.2017.07.008

16 Wang, M. et al. Age-specific associations of RBC folate and several serum folate forms with obesity risk: NHANES 2011-2018. Front Nutr 12, 1547844 (2025). 10.3389/fnut.2025.1547844

17 Yu, X. et al. Folate supplementation modifies CCAAT/enhancer-binding protein α methylation to mediate differentiation of preadipocytes in chickens. Poult Sci 93, 2596–2603 (2014). 10.3382/ps.2014-04027

18 Williams, J. et al. Atlas of one-carbon metabolism in conventional and germ-free mice reveals folate as a key determinant of biochemical pathways. Nat Metab 8, 924–940 (2026). 10.1038/s42255-026-01489-w

19 Wunderling, K., Zurkovic, J., Zink, F., Kuerschner, L. & Thiele, C. Triglyceride cycling enables modification of stored fatty acids. Nat Metab 5, 699–709 (2023). 10.1038/s42255-023-00769-z

20 Hilgendorf, K. I. et al. Omega-3 Fatty Acids Activate Ciliary FFAR4 to Control Adipogenesis. Cell 179, 1289–1305.e1221 (2019). 10.1016/j.cell.2019.11.005

21 Lee, Y., Vousden, K. H. & Hennequart, M. Cycling back to folate metabolism in cancer. Nat Cancer 5, 701–715 (2024). 10.1038/s43018-024-00739-8

22 Genestier, L., Paillot, R., Quemeneur, L., Izeradjene, K. & Revillard, J. P. Mechanisms of action of methotrexate. Immunopharmacology 47, 247–257 (2000). 10.1016/s0162-3109(00)00189-2

23 Zhou, Y. et al. Metascape provides a biologist-oriented resource for the analysis of systems-level datasets. Nat Commun 10, 1523 (2019). 10.1038/s41467-019-09234-6

24 Grabner, G. F., Xie, H., Schweiger, M. & Zechner, R. Lipolysis: cellular mechanisms for lipid mobilization from fat stores. Nat Metab 3, 1445–1465 (2021). 10.1038/s42255-021-00493-6

25 Houten, S. M., Violante, S., Ventura, F. V. & Wanders, R. J. The Biochemistry and Physiology of Mitochondrial Fatty Acid β-Oxidation and Its Genetic Disorders. Annu Rev Physiol 78, 23–44 (2016). 10.1146/annurev-physiol-021115-105045

26 Ghaben, A. L. & Scherer, P. E. Adipogenesis and metabolic health. Nat Rev Mol Cell Biol 20, 242–258 (2019). 10.1038/s41580-018-0093-z

27 Schweiger, M. et al. Adipose triglyceride lipase and hormone-sensitive lipase are the major enzymes in adipose tissue triacylglycerol catabolism. J Biol Chem 281, 40236–40241 (2006). 10.1074/jbc.M608048200

28 Bates, R., Huang, W. & Cao, L. Adipose Tissue: An Emerging Target for Adeno-associated Viral Vectors. Mol Ther Methods Clin Dev 19, 236–249 (2020). 10.1016/j.omtm.2020.09.009

29 Frühbeck, G., Catalán, V., Rodríguez, A. & Gómez-Ambrosi, J. Adiponectin-leptin ratio: A promising index to estimate adipose tissue dysfunction. Relation with obesity-associated cardiometabolic risk. Adipocyte 7, 57–62 (2018). 10.1080/21623945.2017.1402151

30 Crider, K. S., Yang, T. P., Berry, R. J. & Bailey, L. B. Folate and DNA methylation: a review of molecular mechanisms and the evidence for folate’s role. Adv Nutr 3, 21–38 (2012). 10.3945/an.111.000992

31 Duthie, S. J. & Hawdon, A. DNA instability (strand breakage, uracil misincorporation, and defective repair) is increased by folic acid depletion in human lymphocytes in vitro. FASEB J 12, 1491–1497 (1998).

32 Wasson, G. R. et al. Global DNA and p53 region-specific hypomethylation in human colonic cells is induced by folate depletion and reversed by folate supplementation. J Nutr 136, 2748–2753 (2006). 10.1093/jn/136.11.2748

33 Van Vranken, J. G., Li, J., Mitchell, D. C., Navarrete-Perea, J. & Gygi, S. P. Assessing target engagement using proteome-wide solvent shift assays. Elife 10 (2021). 10.7554/eLife.70784

34 Lowe, K. E. et al. Regulation of folate and one-carbon metabolism in mammalian cells. II. Effect of folylpoly-gamma-glutamate synthetase substrate specificity and level on folate metabolism and folylpoly-gamma-glutamate specificity of metabolic cycles of one-carbon metabolism. J Biol Chem 268, 21665–21673 (1993).

35 Shane, B. Folylpolyglutamate synthesis and role in the regulation of one-carbon metabolism. Vitam Horm 45, 263–335 (1989). 10.1016/s0083-6729(08)60397-0

36. Neurosnap Inc. (2022) Neurosnap: An online platform for computational biology and chemistry. Available at: https://neurosnap.ai/.

37 Paudel, R., Fusi, L. & Schmidt, M. The MEK5/ERK5 Pathway in Health and Disease. Int J Mol Sci 22 (2021). 10.3390/ijms22147594

38 Zhu, H. et al. Role of extracellular signal-regulated kinase 5 in adipocyte signaling. J Biol Chem 289, 6311–6322 (2014). 10.1074/jbc.M113.506584

39 Duerre, D. J. et al. Haem biosynthesis regulates BCAA catabolism and thermogenesis in brown adipose tissue. Nat Metab 7, 1018–1033 (2025). 10.1038/s42255-025-01253-6

40 Emont, M. P. et al. A single-cell atlas of human and mouse white adipose tissue. Nature 603, 926–933 (2022). 10.1038/s41586-022-04518-2

41 Efthymiou, V. et al. Single-Nucleus Analysis of Human White Adipose Tissue Reveals Adipocyte Subsets with Distinct Metabolic Profiles. bioRxiv (2025). 10.1101/2025.09.14.673351

42 Zhong, J. et al. adiposetissue.org: A knowledge portal integrating clinical and experimental data from human adipose tissue. Cell Metab 37, 566–569 (2025). 10.1016/j.cmet.2025.01.012

43 Bailey, L. B. et al. Biomarkers of Nutrition for Development-Folate Review. J Nutr 145, 1636S–1680S (2015). 10.3945/jn.114.206599

44 Raz, S., Stark, M. & Assaraf, Y. G. Folylpoly-γ-glutamate synthetase: A key determinant of folate homeostasis and antifolate resistance in cancer. Drug Resist Updat 28, 43–64 (2016). 10.1016/j.drup.2016.06.004

45 Srivastava, A. C. et al. Elimination of human folypolyglutamate synthetase alters programming and plasticity of somatic cells. FASEB J 33, 13747–13761 (2019). 10.1096/fj.201901721R

46 Marques, C. et al. Methotrexate enhances 3T3-L1 adipocytes hypertrophy. Cell Biol Toxicol 29, 293–302 (2013). 10.1007/s10565-013-9255-0

47 Perumal, N. L. et al. BIX02189 Suppresses Adipogenesis and Lipid Accumulation Through Inhibition of MEK5-STAT3/STAT5 Signaling and Activation of AMPK in Adipocytes and Zebrafish. Int J Mol Sci 27 (2026). 10.3390/ijms27146468

48 Drew, B. A., Burow, M. E. & Beckman, B. S. MEK5/ERK5 pathway: the first fifteen years. Biochim Biophys Acta 1825, 37–48 (2012). 10.1016/j.bbcan.2011.10.002

49 Lu, C. et al. Fine Mapping of the. Front Genet 13, 838685 (2022). 10.3389/fgene.2022.838685

50 Clasquin, M. F., Melamud, E. & Rabinowitz, J. D. LC-MS data processing with MAVEN: a metabolomic analysis and visualization engine. Curr Protoc Bioinformatics Chapter 14, Unit14.11 (2012). 10.1002/0471250953.bi1411s37

51 Counter, C. M. et al. Dissociation among in vitro telomerase activity, telomere maintenance, and cellular immortalization. Proc Natl Acad Sci U S A 95, 14723–14728 (1998). 10.1073/pnas.95.25.14723

52 Lee, M. J., Wu, Y. & Fried, S. K. A modified protocol to maximize differentiation of human preadipocytes and improve metabolic phenotypes. Obesity (Silver Spring) 20, 2334–2340 (2012). 10.1038/oby.2012.116

53 Tchkonia, T. et al. Fat depot-specific characteristics are retained in strains derived from single human preadipocytes. Diabetes 55, 2571–2578 (2006). 10.2337/db06-0540

54 Li, J. et al. TMTpro-18plex: The Expanded and Complete Set of TMTpro Reagents for Sample Multiplexing. J Proteome Res 20, 2964–2972 (2021). 10.1021/acs.jproteome.1c00168

55 Rad, R. et al. Improved Monoisotopic Mass Estimation for Deeper Proteome Coverage. J Proteome Res 20, 591–598 (2021). 10.1021/acs.jproteome.0c00563

56 Li, J. et al. TMTpro reagents: a set of isobaric labeling mass tags enables simultaneous proteome-wide measurements across 16 samples. Nat Methods 17, 399–404 (2020). 10.1038/s41592-020-0781-4

57 Huttlin, E. L. et al. A tissue-specific atlas of mouse protein phosphorylation and expression. Cell 143, 1174–1189 (2010). 10.1016/j.cell.2010.12.001

58 Savitski, M. M., Wilhelm, M., Hahne, H., Kuster, B. & Bantscheff, M. A Scalable Approach for Protein False Discovery Rate Estimation in Large Proteomic Data Sets. Mol Cell Proteomics 14, 2394–2404 (2015). 10.1074/mcp.M114.046995

59 Brady, D. C. et al. Copper is required for oncogenic BRAF signalling and tumorigenesis. Nature 509, 492–496 (2014). 10.1038/nature13180

